# A benchmark for large language models in bioinformatics

**DOI:** 10.1101/2023.12.19.572483

**Authors:** Varuni Sarwal, Gaia Andreoletti, Viorel Munteanu, Ariel Suhodolschi, Dumitru Ciorba, Viorel Bostan, Mihai Dimian, Eleazar Eskin, Wei Wang, Serghei Mangul

## Abstract

The rapid advancements in artificial intelligence, particularly in Large Language Models (LLMs) such as GPT-4, Gemini, and LLaMA, have opened new avenues for computational biology and bioinformatics. We report the development of BioLLMBench, a novel framework designed to evaluate LLMs in bioinformatics tasks. This study assessed GPT-4, Gemini, and LLaMA through 2,160 experimental runs, focusing on 24 distinct tasks across six key areas: domain expertise, mathematical problem-solving, coding proficiency, data visualization, research paper summarization, and machine learning model development. Tasks ranged from fundamental to expert-level challenges, and each area was evaluated using seven specific metrics. A Contextual Response Variability Analysis was implemented to understand how model responses varied under different conditions. Results showed diverse performance: GPT-4 led in most tasks, achieving a 91.3% proficiency in domain knowledge, while Gemini excelled in mathematical problem-solving with a 97.5% proficiency score. GPT-4 also outperformed in machine learning model development, though Gemini and LLaMA struggled to generate executable code. All models faced challenges in research paper summarization, scoring below 40% using the ROUGE metric. Model performance variance increased when using a new chat window, though average scores remained similar. The study also discusses the limitations and potential misuse risks of these models in bioinformatics.

## Introduction

In the past decade, there have been rapid advancements in high-throughput leading to a wealth of genomic, transcriptomic, and proteomic datasets, which has fueled a need for innovative computational approaches to extract insights from complex biological datasets. While dataset interpretation is central to understanding intricate biological phenomena ^1,2^, the extraction of meaningful, novel insights presents a challenging endeavor due to their inherent complexity and volume of these datasets. Recent advancements in artificial intelligence ^3^ and particularly in the development of large language models (LLMs), offer a promising direction in the pursuit of efficient and robust multimodal data interpretation in bioinformatics^4,5^. LLMs, such as GPT-4^6^ (the fourth iteration of the Generative Pre-trained Transformer model (GPT-4), trained by OpenAI) and Gemini^7,8^ (Binary Augmented Retro-Framing, a conversational generative artificial intelligence chatbot based on Pathways Language Model (PaLM2), developed by Google), have demonstrated remarkable capabilities in a wide array of natural language processing tasks^9^.

These models are built upon state-of-the-art deep learning architectures, trained on massive amounts of text data to simulate human conversations, and promise to generate coherent and contextually relevant responses across a range of scientific domains^10^ . LLMs, with their ability to learn patterns, semantic relationships, and hidden structures in unstructured text data, offer a new perspective to assist bioinformatics research. They hold promise for tasks such as gene expression analysis, variant interpretation, protein folding prediction, and drug discovery.^11,12^ Previous studies have explored practical aspects of harnessing the GPT-4’s capabilities through prompt design, as well as potential use cases where GPT-4 can assist bioinformatics analysis^13–15^. Another recent study has discussed the potential risks of using GPT-4 for bioinformatics tasks, such as privacy risks when used for medical EHR-based datasets^15^. In addition, in a recent paper, authors present DNAGPT, a generalized pretrained tool for DNA sequence analysis^16^. Notwithstanding the immense potential of LLMs, the extent to which these models can contribute to bioinformatics research and education remains largely unexplored^2,17^.

The accuracy and reliability of the LLMs and ease of usage by non-experts in the fields such as genetics^18^ or microbiology^19^ remains an open question. Additionally, there is a crucial need for a study to acknowledge and address the associated risks and limitations to ensure responsible and ethical use in this field^20,21^. Therefore, it is essential to develop an updated benchmark to evaluate the research progress of LLMs in solving bioinformatics research based problems.

Bioinfo-Bench^22^ was recently introduced as a simple yet broad evaluation suite to assess an LLM’s academic bioinformatics knowledge and basic analytical skills. It focuses on testing foundational knowledge acquisition and reasoning in bioinformatics rather than end-to-end workflows. The intended use case is to gauge how well models retain and apply domain knowledge (for instance, answering textbook-style questions or simple data interpretation) as a baseline for bioinformatics expertise in LLMs. Bioinfo-Bench’s dataset provides a broad knowledge test suite in bioinformatics, but with limited depth in each sub-area and no coding data.

In this paper, we present BioLLMBench, a novel computational framework and a scoring metric scheme for evaluating LLMs in bioinformatics. Using BioLLMBench, we conduct a comprehensive evaluation of LLMs by assigning them daily tasks encountered by bioinformatics researchers. These tasks consisted of six key areas, namely domain-specific knowledge, coding, visualization, machine learning (ML), mathematical problem-solving, and research paper summarization. The tasks also span across varying levels of complexity, ranging from fundamental concepts to expert-level challenges. We evaluate over 2,160 experimental runs of the three most widely using models, GPT-4, Gemini and LLaMA, focusing on 24 distinct tasks within the field of bioinformatics. These were hand-crafted to mimic real bioinformatics challenges To enhance our understanding of model responses under varying conditions, we implemented a Contextual Response Variability Analysis. This involved a dual-phase process where each task was evaluated through 10 runs maintaining a consistent search window and query, followed by another 10 runs in a new search window for each model. This methodical approach was crucial for investigating how changes in the immediate contextual environment (same versus new chat window) influence the variability and consistency of the models’ responses across different levels of task complexity and domains.

## Methods

### Model Selection

Four models were chosen for evaluation:

- Gemini Pro 1.0: A multimodal model developed by Google, optimized for conversational AI tasks through Vertex AI APIs.
- GPT-4: OpenAI’s advanced generative model with multimodal capabilities and a large context window of up to 128,000 tokens.
- LLaMA 2 7B: Meta’s compact language model optimized for dialogue-based tasks with fine-tuning on human annotations.
- PaLM: Google’s Pathways Language Model designed for large-scale natural language processing tasks.

### Designing The Experiments

The panel covered a range of topics specific to six key areas of emphasis within bioinformatics that directly relate to the daily challenges and tasks commonly faced by individuals within the bioinformatics field. These areas encompass domain expertise, mathematical problem-solving, coding proficiency, data visualization, summarizing research papers, and developing machine learning models. The questions were generated over 4 levels of difficulty, from easy, moderate, high, to expert level. Domain knowledge tasks were designed to test the LLM’s understanding of bioinformatics concepts and principles. Answering these questions required an in-depth domain knowledge of different fields within bioinformatics. This included questions about genomic annotation, sequence alignment, genome assembly, and more (Table 1). Coding tasks were designed to evaluate the LLM’s ability to generate correct and efficient code for common bioinformatics problems, such as parsing a FASTA file or calculating GC content. The LLMs produced snippets of code, which were then run to test correctness. Visualization tasks assessed the LLM’s understanding of how to generate and interpret common bioinformatics plots and graphs. Visualization problems required synthetically generating a biological dataset, and then generating code in Python that could visualize the data. The code was run to test the correctness and completeness of the visualizations. In order to test the mathematical accuracy of the LLMs, we generated questions related to bioinformatics that required a mathematical calculation to reach the answer and had only one correct response. The answers produced by the LLMs were compared to a human-generated gold standard. For machine learning, we used the University of California, Irvine (UCI) heart disease binary classification dataset. The UCI heart classification dataset is a multivariate EHR dataset composed of 14 features attributes, namely age, sex, chest pain type, resting blood pressure, serum cholesterol, fasting blood sugar, resting electrocardiographic results, maximum heart rate achieved, exercise-induced angina, old peak ST depression induced by exercise relative to rest, the slope of the peak exercise ST segment, number of major vessels and Thalassemia. We used the UCI dataset since it is a publicly available healthcare-ML dataset with a well-established benchmark. While we provided a csv file containing the features and outcome to GPT-4, we asked LLaMA and Gemini models to download the data themselves. Then, our prompt instructed the models to perform data preprocessing, feature selection and model training for building a classification model given the dataset. In case of an error message, we provided the error generated to the LLM as input, and asked it to debug. We tested the alternate code that the LLM provided in a series of iterations. If the model sustained the same error for more than 10 iterations, a score of 0 was provided. The final accuracy at the end of the debugging process was reported. Reading research papers is an important task for bioinformatics researchers, so we devised a research paper summarization challenge in which we asked the LLM’s to summarize the top 10 most cited papers in bioinformatics of all time (https://github.com/manjunath5496/The-10-most-cited-Bioinformatics-papers-of-all-time).

**Table 1:** A summary of metrics used for the LLM response evaluation.

|  |  |
| --- | --- |
| Domain knowledge |  |
| Metric | Explanation |
| Accuracy | Whether the information provided is correct and up to date. The evaluation was based on known facts and standard practices in the domain. Points were deduced for any inaccuracy or misleading statements. |
| Completeness | Whether the response covers all aspects of the topic – including definitions, applications and limitations. |
| Relevance | The degree to which the content of the response is pertinent to the question asked. |
| Depth and Details | The level of detail and thoroughness of the explanation provided. |
| Clarity | How easily understandable the response is – the use of language, the complexity of terms used, explanation of complex concepts. |
| Organization | The structure and layout of the response – whether it had a logical flow, and the use of bullet points for better readability. |
| Conciseness | The brevity of the response in relation to the amount of information provided – avoiding unnecessary repetitions/formulations, while still being complete and informative. |
| Code writing |  |
| Metric | Explanation |
| Readability and Style | How easy it is to read and understand the code and the accompanying explanations/comments. Factors include code formatting, use of comments, and the overall presentation of the code and explanation. |
| Correctness | Evaluates whether the code correctly performs the task. It assesses the accuracy of the code's logic and output, ensuring that the solution aligns with standard practices/formats and produces the expected results. |
| Efficiency | How well the code performs computationally, in term of resource usage (memory and processing time). |
| Simplicity | Complexity of the code: based on this metric we favor solutions that are straightforward and avoid unnecessary complexity, thus making the code more maintainable and easier to understand. |
| Error Handling | Evaluates whether the code includes measures to manage exceptions and incorrect inputs. |
| Code Example | Evaluates whether the code is accompanied by a good example to accomplish the task, and whether it includes important aspects like functions, logical flow, and integration of necessary libraries. |
| Clear Input and Output | Evaluates how clearly the code defines expected input and output in terms of formats and file types. |
| Visualization |  |
| Metric | Explanation |
| Clarity and Simplicity | Evaluates how easy it is to understand the response (clear language, avoiding jargon, easy-to-follow manner). |
| Instructions | Evaluates whether the code provides a clear, logical, and sequential guide. |
| Data Preparation | Evaluates how well the response guides the reader in preparing and understanding the data for the task. |
| Axis Labels and Legend | Evaluates if there are instructions or guidance on labeling the axes of the visualizations and using legends effectively to clarify information. |
| Color usage | The use of graphical grammar. |
| Example providing | Evaluates if the response is based on code snippets or examples that could be used to directly implement the visualizations. |
| Interpretation | Evaluates how well the response provides an interpretation of results for visualization. |
| Software and Tools | Evaluates if there are recommendations for specific software or tools (packages and libraries) that can be used for creating the plots. |

We excluded the abstract of the research paper from the input, and used the abstract as the gold standard summary. All of the papers were input via the chat interface that was provided by developer except for LLaMA-2, where the Graphical User Interface was used (https://github.com/oobabooga/text-generation-webui) running on a public URL in the browser as a base for input of data. The LLM’s were prompted to summarize the article.

### Running The Large Language Models

Questions from each category were input to the GUI-based interface for each of the models. We chose to run the models using the GUI interface instead of performing an API call since many bioinformatics researchers are unfamiliar with the command line interface. The default parameters were used. Each question was prompted to each of the LLMs 20 times, 10 times in the same chat window and 10 times in a different chat window. Finally, the responses were saved and evaluated with metrics specific to each task. Gemini generates 3 answer templates at the same time, so we considered the first template of each response set. Gemini would also produce the following message at certain instances: “That’s all Gemini can answer right now.

It’s experimental. Try back later”. During instances where this phenomenon occurred, we waited about 15 (+-5) seconds before trying again. We observed that extending the wait time to 3–60 seconds minimizes the number of “try back later” messages. LLaMA was the only model setting of which was altered for the possibility to obtain full responses without cutting the generation at the beginning. LLaMA was run using Google Colab via accessing and downloading the model of LLaMA 2 7b version on huggingface. Then, using Graphical User Interface running on a public

URL in the bowser as a base for input of data. All of the model settings were left default except if max_new_tokens parameters were set to 4096. Generation attempts parameters were set to 10 and length parameters were set to 4864. This combination of parameters was the best for obtaining the desired results. GPT-4 was the only model for which a paid version was available for this study, while Gemini and LLaMA were free. Since the performance of LLM’s is rapidly evolving, it is important to note that the time frame of these experiments is from June-September, 2023.

### Performance Evaluation

We designed a comprehensive evaluation framework to evaluate the response of LLMs, tailor made for each of the tasks. For each of these metrics, the response was analyzed by experts and scored from 1 to 10 with a full score of 10 corresponding to a correct and fully complete response. The scores reflect the degree to which the response met each metric’s criteria. We defined the Consolidated Proficiency Index (CPI), representing the average normalized performance of each LLM across seven task specific metrics for each key area. Each sub-metric is normalized to a [0-10] scale, with 10 indicating the best possible performance. The final score is the mean of these normalized values, providing a comprehensive measure of the LLM’s overall capabilities in the field. This approach ensured a balanced assessment of tasks, considering various aspects of the provided responses. For math-based questions, the scoring was binary, meaning the scores were either 0 or 1, based on the correctness of the response. Details of the evaluation framework can be found in Table 2. For the coding challenge, we considered the accuracy produced by the model. For the research paper summarization task, we used the paper abstract as the gold standard summary. Then, we computed the rouge score to measure the degree of similarity between the abstract and the LLM generated summary.

**Table 2:** Results from the machine learning challenge.

| Model | Accuracy | Error Handling | One hot encoding | # Iterations | Model selection | Data Imputation |
| --- | --- | --- | --- | --- | --- | --- |
| GPT-4 | 63.32% | 9.8 | 10 | 1.6 | Random Forest: 80%, Logistic Regression: 10%, Gradient Boosting Trees: 10% | Mean: 60%<br>Median: 40% |
| Gemini | 0.00% | 0.8 | 0 | 10 | Random Forest: 70%, Decision Tree: 10%, Logistic Regression: 10%, N/A: 10% | Mean: 10%<br>N/A: 90% |
| LLaMA | 0.00% | 0.0 | 1.25 | 10 | Logistic Regression: | N/A: 100% |
|  |  |  |  |  | 70%, N/A:<br>30% |  |

The average across model accuracy is displayed. Error handling and One hot encoding skills are on a scale of 1 to 10. When the number of iterations required was more than 10, a score of 10 was assigned. The distribution of the classification model used by the LLM (model selection) and data imputation methods for missing data chosen by the models are summarized.

## Results

### Evaluating Large Language Models in Bioinformatics

BioLLMBench was developed to evaluate LLM performance on bioinformatics tasks using GPT- 4, Gemini, and LLaMA. Six key areas were evaluated using seven task-specific metrics, and a Consolidated Proficiency Index (CPI) was used to measure overall LLM capability (Table 1-2). We applied this framework across 24 distinct tasks (see Methods). We carefully hand-selected these tasks to cover a broad spectrum of bioinformatics concepts, ensuring a comprehensive coverage of the field. The tasks also span across varying levels of complexity, ranging from fundamental concepts to expert-level challenges (Tables 1 and 2). In total, we analyzed the results from 2,160 experimental runs, and for each task, we performed 20 runs per model, with 10 runs of asking the model the same question in the same search window, and 10 runs using a new search window. This was done to analyze if there were any differences between following both processes, and advise users on which method to use based on their domain expertise and downstream application. Once we received the responses generated by the LLM for both settings, we used our evaluation framework to analyze the quality of each response.

First, we evaluated the domain knowledge performance of the language models. For this task, we included questions that assessed the model’s understanding of biological and bioinformatics fundamental concepts. Most specifically, we included questions that involved defining common bioinformatics concepts, and describing the difference between related topics (Table S1). For example, we asked the models to define ‘Genome Annotation’ and explain its importance. Each of the models provided a descriptive textbook based response of genome annotation. We evaluated the model responses based on the accuracy, completeness, conciseness, and clarity of the response. In addition, we also considered how relevant the response was to the question asked, and the amount of details and depth provided. Finally, we also evaluated the organization of the structure of the responses by each of the models. Next, we tested the ability of the models in generating code for scientific visualization. For example, for a given set of p-values from a GWAS study, we asked the models to describe how to visualize the p-values in a Manhattan plot. We ran the code and evaluated the quality of the code in terms of the clarity and simplicity of the code, and the figure attributes such as the axis labeling and color usage.

We also evaluate whether the model provides a code snippet as an example, the data preparation process (if required), the logical sequence of instructions provided, and how well the results can be interpreted through the visualization.

Bioinformatics analysis frequently requires code for data analysis, and we generated a set of coding questions to test the ability of LLMs to write compilable code. For example, we asked the models to write a Python function to calculate the Hamming distance between two DNA sequences. This code was then compiled and checked for errors, and run to test the output. We evaluated the code in terms of readability and style, correctness, efficiency, and simplicity, the error handling ability, whether the model provided a code example, and the clarity of the input and output data. In order to evaluate the problem-solving skills of the LLMs in bioinformatics tasks, we created a set of bioinformatics questions that required mathematical computation and had only one correct answer. For example, we asked the model to compute the percentage of bases in a DNA sequence, the Bonferroni corrected p-value for GWAS, and count the number of reads generated in sequencing. The LLMs were prompted to produce a single numerical response, which was then compared to a human-generated gold standard. Scoring was binary, with a score of one assigned for the correct answer, and a zero otherwise.

Reading and summarizing research papers is an important skill in research fields including but not limited to bioinformatics. To assess model capabilities, we tested the performance of LLMs to summarize scientific texts. For this challenge, we provided the top 10 most cited bioinformatics papers to the 3 LLMs, and asked them to generate a summary. Using the abstract as the ground truth summary, we computed the rouge score to measure the semantic similarity between the abstract and the summary generated by the LLM.

The machine learning challenge was designed to test if the LLMs could develop an end-to-end classification model. We used the popular UCI heart disease dataset^23^, which is a multivariate EHR dataset composed of 14 feature attributes. We asked the models to perform data preprocessing, feature selection, data imputation, and model selection. Finally, we asked the model to report the classification accuracy. We scored the model based on several criteria: the number of iterations required to produce working code, the accuracy of the binary classification, the simplicity of error handling, its ability to one-hot encode categorical variables, and the data imputation method it employed.

### Language Models Excelled In Information Retrieval And Comparison For Bioinformatics Domain Knowledge Tasks

We evaluated the performance of GPT-4, Gemini and LLaMA on information retrieval tasks in the bioinformatics domain. These included tasks on concept definitions and differentiation between related concepts. We observed all three models to have a CPI in this category, with GPT-4 emerging as the top-performing model, with a CPI of 9.13, followed by LLaMA and Gemini (Figure 1a, Figure 2a). Our observations indicate that all models consistently performed well in tasks involving an understanding of bioinformatics concepts across most complexity levels, and accurately answered questions related to genomic annotation, sequence alignment, and genome assembly, among others, even at the expert level. The models were also excellent at differentiating between concepts. For example, when asked to differentiate various sequencing protocols, Gemini effectively provided baseline definitions for each method. It then produced a table summarizing the key differences across five key features. On the other hand, while LLaMA provided the key differences, it occasionally included irrelevant information, such as potential future improvements in whole genome and exome sequencing. While GPT-4 had the highest scores overall, there were instances in our experiments where GPT-4 tended to oversimplify concepts, providing very concise responses (Figure 2a, Supplementary Materials). Conversely, LLaMA provided more comprehensive responses with higher depth and details, as well as completeness (Figure 2a), while it had a lower performance on the accuracy, clarity, organization and relevance of the response to the topic. While Gemini had the lowest overall score across all metrics, it uniquely provided the capability to search the prompt on Google and search information related to the response. This could be particularly useful for fact-checking and follow-up searchers. Looking closely, we observe that most of Gemini’s individual scores were high, except for one iteration, where it responded to the query with “I’m a text-based AI and can’t assist with that”. Gemini also frequently reached the maximum question limit, and would subsequently decline to provide a response. Multiple prompting can be useful in such scenarios (Figure 4).

**Figure 1:**
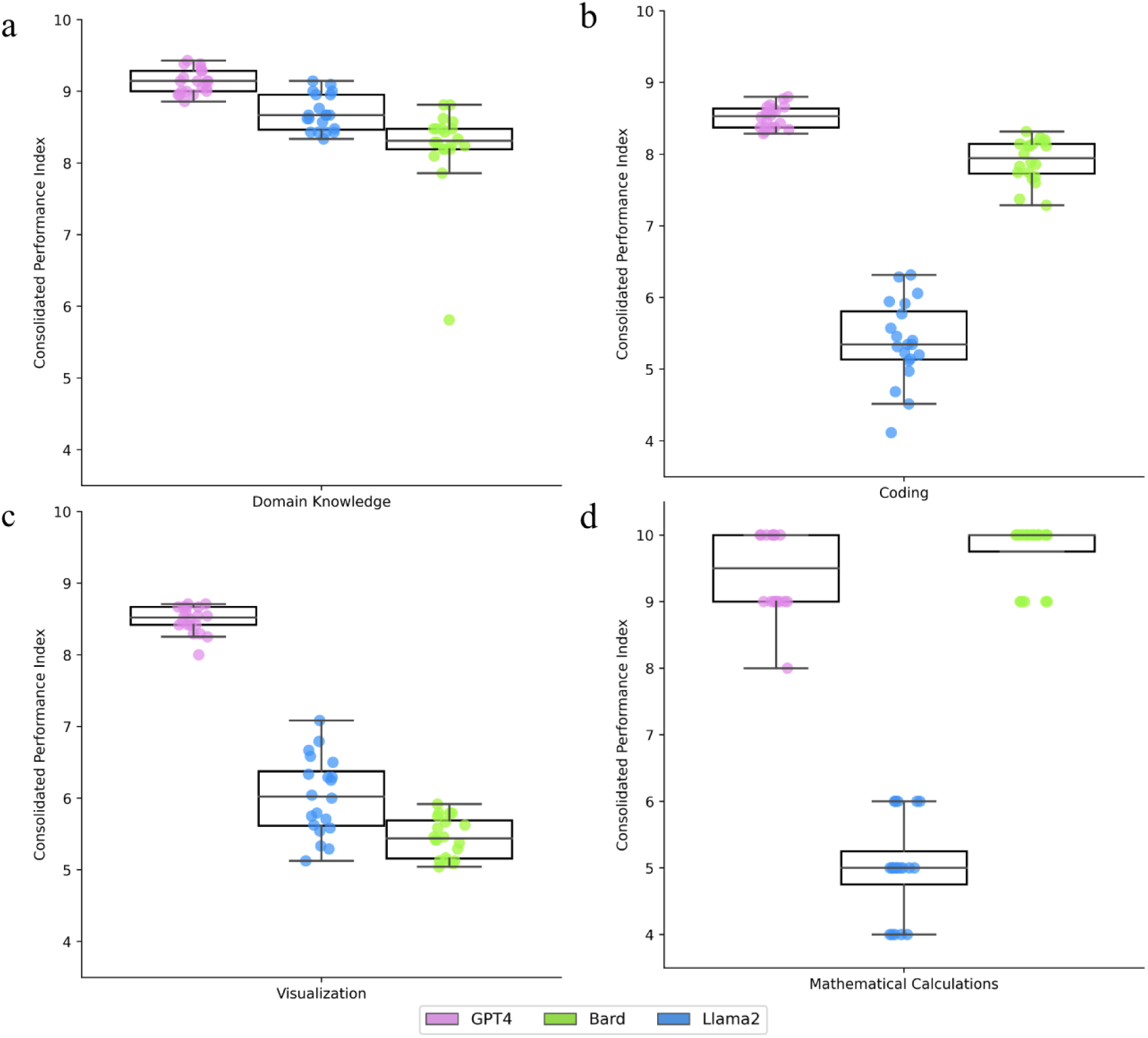
LLMs performance across four different knowledge metrics. A box plot chart displaying the performance of GPT-4 vs Gemini on common bioinformatics tasks in the following categories (a) Domain knowledge tasks: Each point corresponds to a unique run for the same chat and new chat window. The CPI is the average score over 7 metrics: Accuracy, Completeness, Relevance, Depth and Details, Clarity, Organization and Conciseness (b) Coding tasks: Each point corresponds to a unique run for the same chat and new chat window. The CPI is th e average score over 8 metrics: Readability and Style, Correctness, Efficiency, Simplicity, Error Handling, Example provided and Clear Input and Output (IO) provided or not. (c) Visualization: Each point corresponds to a unique run for the same chat and new chat window. The CPI is the average score over 8 metrics: Data, Instructions, Clarity and Simplicity, Software, Interpretation, Example, Color, and Axis Labels and Legend

**Figure 2:**
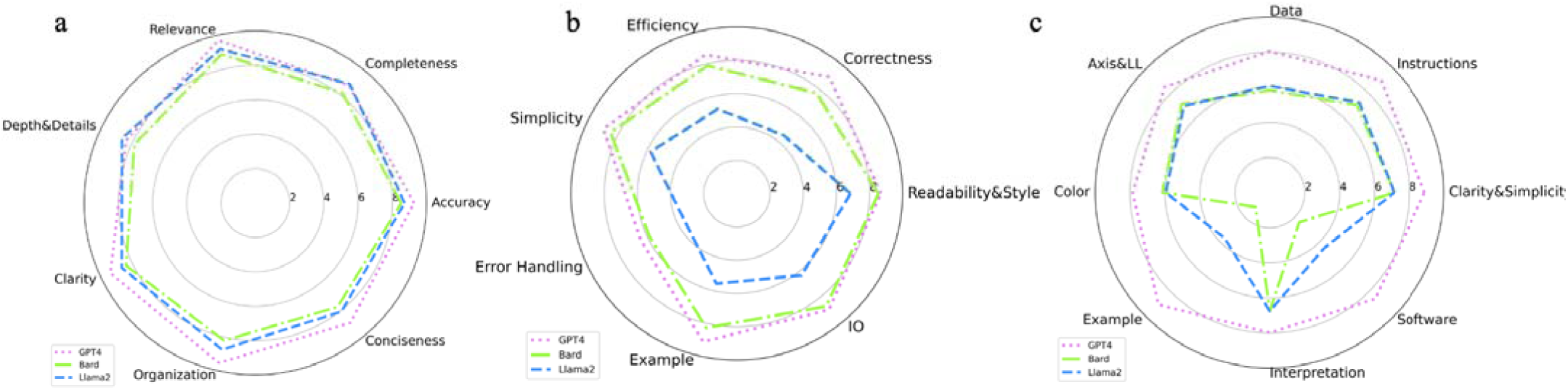
LLM performance parametrized by evaluation metrics specific to each task(a) LLMs performance on coding tasks. The average over 20 runs for the same chat and new chat windows for each LLM is provided. We evaluate the models over 7 code-writing metrics: readability and style, correctness, efficiency, simplicity, error handling, example provided and clear input and output (IO) provided or not. (b) LLMs performance on visualization tasks. The average over 20 runs for the same chat and new chat windows for each LLM is provided. We evaluate the model over 8 visualization metrics: data, instructions, clarity and simplicity, software, interpretation, example, color, and axis labels and legend.(c) LLMs performance on domain knowledge tasks. The average over 20 runs for the same chat and new chat windows for each LLM is provided. We evaluate the models over 7 metrics: Accuracy, Completeness, Relevance, Depth and Details, Clarity, Organization and Conciseness.

### Gpt-4 And Gemini Excelled In Bioinformatics Code Generation

We observed a considerable variation in the CPI across models on coding tasks, with both GPT-4 and Gemini demonstrating proficiency in generating code across diverse bioinformatics tasks. While GPT-4 was the top performing model with a mean CPI of 8.5 followed closely by PaLM2, with a median CPI of 7.91, LLaMA was far behind, with a median CPI of 5.4 (Figure 1b).

Although GPT-4 outperformed Gemini across all of the metrics, the difference was the most pronounced in the correctness of the response (Figure 2b). This meant that GPT-4 was better at producing more accurate code, aligning with the standard practices to produce the expected result. On the other hand, the GPT-4 and Gemini’s outputs were the most similar in their readability and style. This meant that both LLMs provided enough comments along with the code, in order to facilitate the users to understand the process. The overall presentation of the code, along with its explanations was similarly of high quality. Although we only asked both models to provide the code, both GPT-4 and Gemini exceeded our expectations by not only providing the requested code but also including example use cases. Furthermore, Gemini consistently supplemented the code examples with the expected output, and an explanation of the code. This can prove to be highly valuable for debugging and validating the correctness of the generated code.

Our observations suggest that language models can be a useful resource to assist researchers in generating code. For example, when tasked with writing a Python function to calculate the Hamming distance between two DNA sequences, GPT-4 first defined the concept of Hamming distance as the number of differences between two DNA strands. It then defined a function to compute the Hamming distance for two sequences. Finally, it provided some sample sequences and instructions on how to run the function with these examples, and the expected results. Additionally, it included an explanation of the code, that can help the user understand the control flow.

### Gpt-4 Provided Functional Code For Visualizations, While Gemini And Llama Excel At Textual Descriptions

We assessed the abilities of GPT-4, Gemini, and LLaMA in generating visualizations for various bioinformatics datasets. Similar to the coding task, we observed a wide variation in the performance across existing models. GPT-4 demonstrated the highest CPI of 8.4 in describing how to generate and interpret common bioinformatics plots and graphs, providing functional code for each task. On the other hand, LLaMA and Gemini only provided textual explanations, and only occasionally provided code (Figure 1c). It is important to note that, at the time of our assessment, neither of the models could produce the visualizations directly ; they only offered code snippets.

Interestingly, while Gemini outperformed LLaMA on general coding tasks, LLaMA was the better performing model on visualization tasks, with mean CPIs of 6.02 and 5.42 respectively.

Furthermore, in contrast to coding and domain knowledge tasks, where the model scores on individual metrics followed the same trend as the overall scores, we observed a notable difference across certain metrics for visualization for Gemini and LLaMA. The difference between LLaMA and Gemini was the most pronounced in terms of whether an example was provided or not, as well as the software used to create the plot (Figure 2c). On the other hand, the two models had similar performance in terms of the interpretation provided by the visualization, the clarity and simplicity of the explanation, and the clarity of the data preparation process. Furthermore, Gemini marginally outperformed LLaMA in terms of the plot characteristics, consisting of axis labeling, legends and color usage.

We observed that GPT-4 can be used as an assistant for generating meaningful end-to-end visualizations. For example, we asked the models a classic coding problem, on how to visualize p-values from a GWAS results in a Manhattan plot. The response of GPT-4 was structured in the following way: first, it provided a text based description of what a Manhattan plot is, and how it can be used to identify regions of the genome containing genetic variants affecting the trait of interest. Then, it proceeds to provide a text based description of how to create a Manhattan plot, right from data organization, coloring chromosomes, plotting, and adding significance threshold lines, to labeling and customizing the plot. Finally, it ended the response by providing an executable snippet in python using matplotlib and pandas Python modules.

### Gemini Emerged As The Top Mathematical Problem Solver, Closely Followed By Gpt-4

We evaluated the performance of GPT-4, Gemini and LLaMA in solving problems that required mathematical computation, each having a unique correct answer (Figure 1d, Table S1).

Interestingly, this was the only task where Gemini outperformed GPT-4, with an accuracy of 97.5%. Notably, while GPT-4 still had a high performance, with a median accuracy of 94.5%, LLaMA was far behind with a median accuracy of only 50%.

We observed LLaMA to perform very poorly on mathematical tasks, providing erroneous answers even for relatively simple questions. For example, we asked the model the following question: “A protein is made of 300 amino acids. How many nucleotides are needed to code for this protein?” We observed highly inconsistent responses over multiple runs of the model, ranging from 14 to 18,000 nucleotides. Even when producing the correct answer consistently, the formatting of the response varied across runs including differences in return data types (int, float) and structure of response (single word versus complete sentence). Furthermore, our findings indicate that the models’ performance was highly influenced by human interactions.

When GPT-4 received feedback that its response was incorrect, it exhibited the tendency to modify its subsequent response, even for initially correct answers. This behavior could potentially be problematic for users without comprehensive domain knowledge.

Finally, we did not observe any correlation between the difficulty of the question and the LLM’s performance. While GPT-4 excelled on both medium and high complexity questions, with near perfect scores for each run, LLaMA achieved scores of zero even for very simple questions, due to incorrect outcomes. Caution should be exercised while using these models for mathematical calculations.

### Gpt-4 Is Able To Build An End-To-End Machine Learning Classification Model

All three LLMs were evaluated on a supervised ML classification task, specifically predicting whether a person had heart disease or not, based on the 14 attributes. Developing a machine learning model end-to-end is a complex task, requiring numerous steps, including data preprocessing, feature selection, model selection and evaluation. While the models could not execute the code themselves, we used the commands provided by the LLMs as command line input, and executed the code ourselves. None of the models were able to develop the model in a zero shot manner, and required human intervention and multiple iterations of prompts to achieve results. The first notable challenge encountered was the nature of our tabular data, which consisted of a mix of continuous and categorical variables. None of the LLM accounted for categorical variables in the data, leading to initial errors. On providing the error message generated as an input to the LLMs, GPT-4 immediately resolved the issue by providing a solution involving one-hot encoding for all categorical data whereas LlaMA and Gemini only encoded the specific column mentioned in the error message, making debugging much slower. The second major challenge was data imputation, where none of the LLMs accounted for missing values in the data, which also required additional prompts for possible solutions for imputation. Finally, the models used a variety of ML models to perform classification. After resolving the error messages, GPT-4 was able to achieve an average classification accuracy of 63% with an average of 1.6 iterations. The highest performance was an accuracy of 89%, with a random forest model chosen with mean imputation. In general, random forest was the most popular model selected by GPT-4, occurring 80% of the times, followed by a logistic regression and gradient boosting model. On the other hand, both Gemini and LLaMA required more than 10 iterations for error handing, and were unable to provide executable code. We observed LLaMA to have a higher lag time while processing the inquiries, and both LLaMA and Gemini had a reduced context length compared to GPT-4, in which the model was not able to contextualize subsequent prompts following the first, if not informed explicitly. Both LLaMA and Gemini also provided incomplete code frequently, requiring the user to prompt for the completion. It is important to note that while GPT-4’s achieved the highest accuracy of 89% and outperformed the other models, it is still lower than the state-of-the-art accuracy of 0.96%^5^.

### Large Language Models’ Encounter Challenges In Effectively Summarizing Research Papers

In order to evaluate the LLMs summarization abilities, we provided the models with the 10 most cited bioinformatics papers to date. We used the paper’s original abstract as the ground truth summary, and computed the rouge scores to measure the quality of summarization. Overall, the model performances were poor, with rouge scores consistently below 40% (Figure 3). GPT-4 was the top performing model overall, followed by Gemini and LLaMA. We observed each of the methods to have its own drawbacks. While GPT-4 provided the highest summarization quality, Gemini exhibited the advantage of providing more detailed summaries compared to GPT-4, offering multiple paragraph options for better understanding. However, it faced a technical limitation of not being able to read papers from biorxiv, although it could access papers from arxiv and perform web scraping. We found no documentation or online support addressing this particular issue. On the other hand, while GPT-4 was able to read papers when provided multiple links referencing related articles in the summary. This is not useful since the summary is expected to contain all the relevant information. On the other hand, Gemini provided code in its summaries, implementing the methods described in the paper, with an example input-output and interpretation. Finally, the summaries provided by LLaMA were of low quality, often omitting key details.

**Figure 3:**
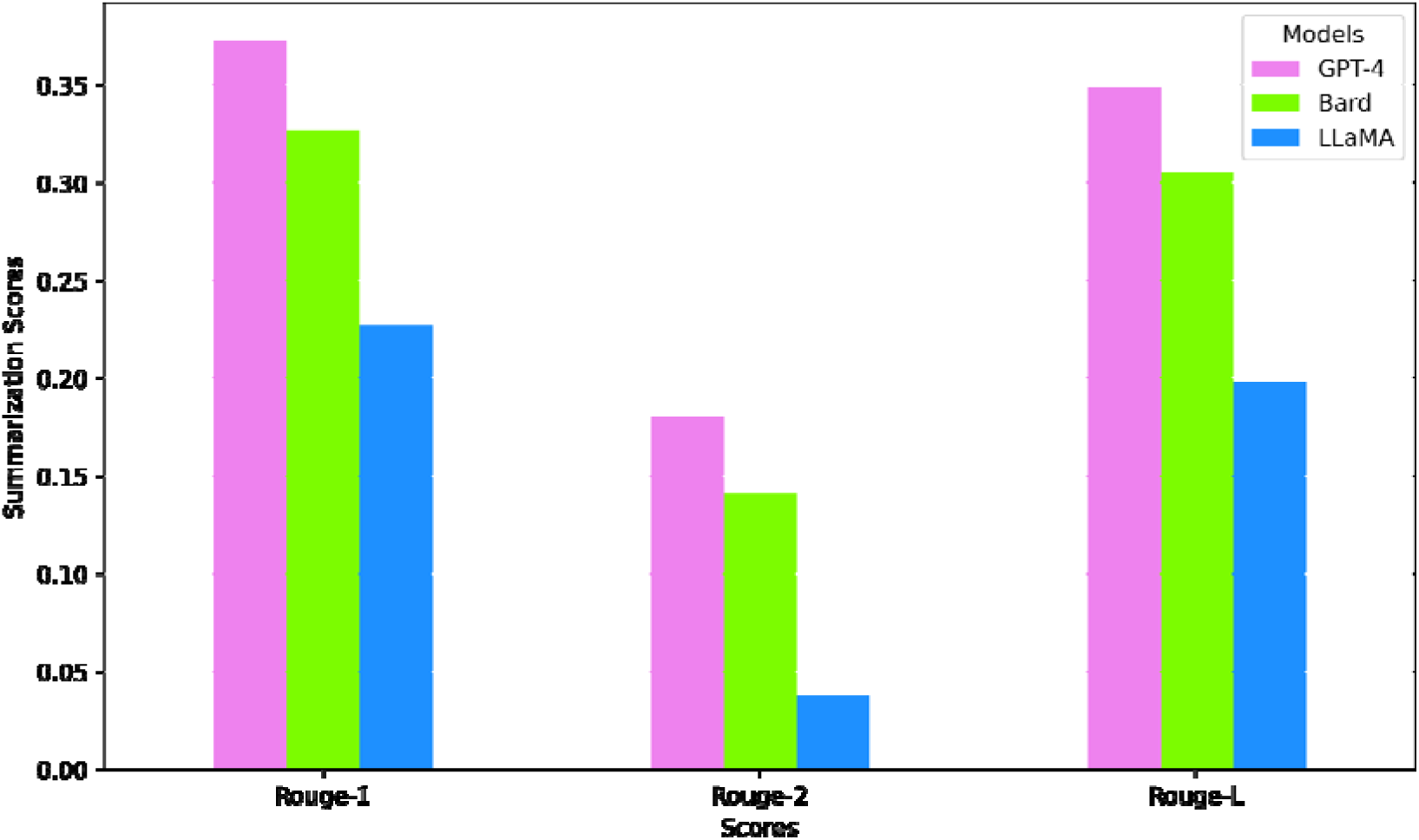
Recall-Oriented Understudy for Gisting Evaluation (ROUGE) scores for research paper summarization for GPT-4, Gemini and LlaMA.

For evaluating the quality of the summarizations we computed the rouge-1, rouge-2 and rouge-L scores. We observed all three models had the highest rouge-L scores, followed by rouge-1 and rouge-2 (Figure 3). This means that the machine-generated summaries were fluent and were able to capture key words from the original text well. A higher rouge-L score indicated that the models were able to capture sentence level structural similarity. Finally, the low rouge-2 score revealed that the machine generated summary was unable to capture the grammatical structure of the abstract, and the ordering of the context. These inconsistencies and low scores raise important concerns regarding the reliability of the summarization output, demanding further investigation and careful consideration from LLM model users.

### **Large Language Models’** Output Variability Increases With New Chat Sessions

In order to analyze the effect of the chat context on the LLMs, we conducted an in-depth investigation into the effects of initiating a new chat versus continuing in an existing one. For each bioinformatics task, we ran the models through 20 rounds, with 10 runs in the same window, and 10 runs using a separate chat window. The average scores across runs were similar for both modes, with scores using an old chat marginally higher for most tasks except visualizations using LLaMA, and coding using GPT4 (Figure 4). On the other hand, we observed a notably higher variance while using a new chat window for most tasks, with the exception of visualization tasks using Gemini, and domain knowledge tasks using GPT4 and LLaMA. Since the average scores were similar, using a new chat window provided extremely high scoring responses in certain iterations while poor responses in others. On the other hand, the same chat responses were fairly consistent, but had an intermediate score. Interestingly, we observed the highest difference of standard deviation for Gemini in domain knowledge questions. This was due to the fact that while most iterations produced extremely high quality responses, one instance of a new chat produced the following response with a zero score: “I’m a text-based AI and can’t assist with that”. On the other hand, LLaMA provided extremely consistent responses for both the same and new chat windows on coding tasks, while the average score for that category was the lowest. These observations suggest that for bioinformatics trainees or users with limited domain expertise, maintaining a consistent chat window could be beneficial for obtaining more stable and reliable model responses. Conversely, for domain experts, the higher variability in new chat windows might present opportunities for producing higher quality, although less predictable, responses from the models.

**Figure 4:**
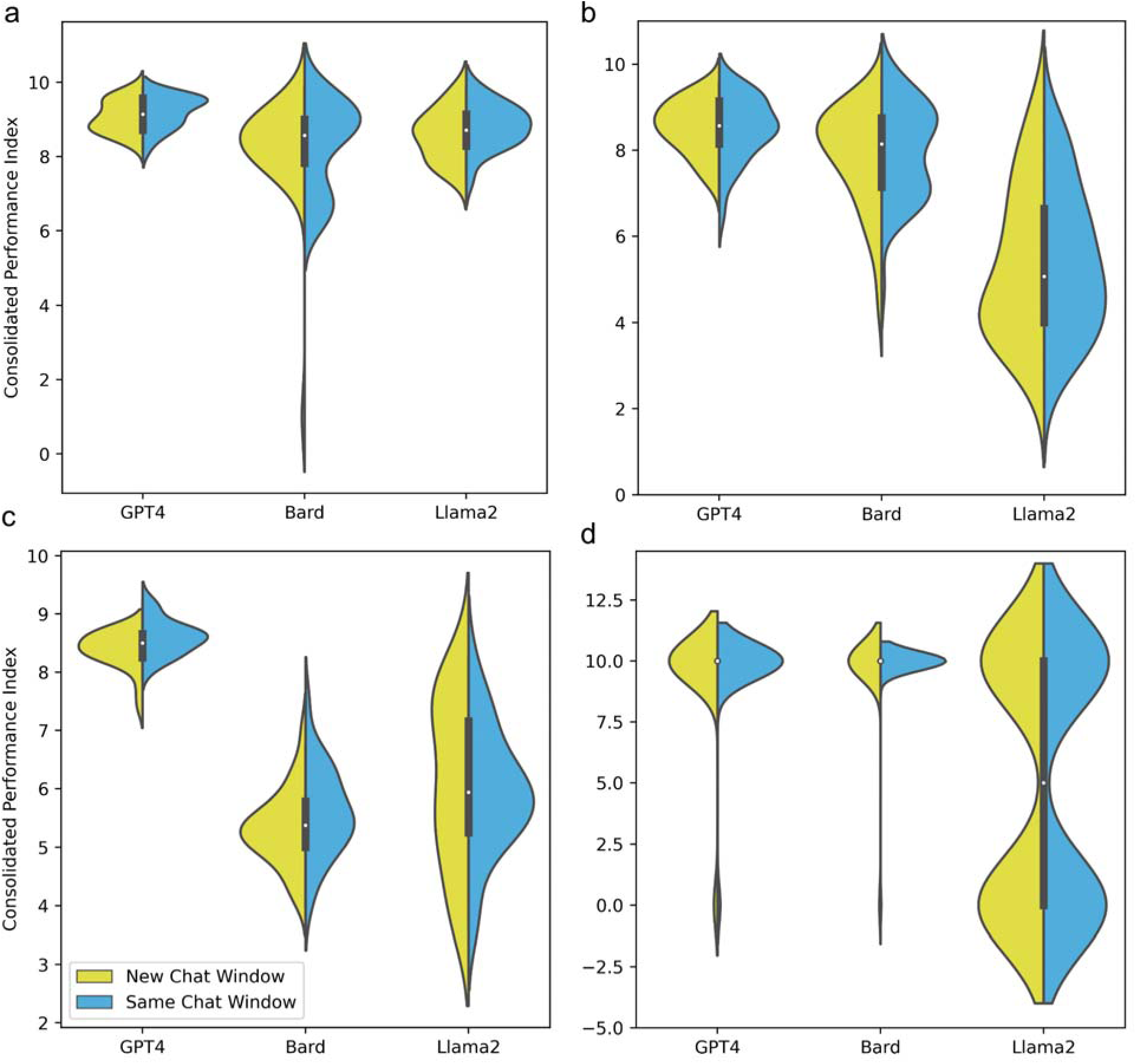
Consolidated Performance Index (CPI) across the same chat (SCH) vs different chat (NCH) for (a) Domain Knowledge with Gemini (b) Coding (c) Visualization and (d) Mathematical Problem Solving

## Discussion

Our study presents BioLLMBench, an assessment framework coupled with a comprehensive scoring metric scheme designed to evaluate the 3 most widely used LLMs, namely GPT-4, Gemini and LLaMA in solving bioinformatics tasks. Among these tools, the GeneTuring^24^ dataset provides an in-depth evaluation of LLMs’ genomic knowledge via a large set of questions, focused solely on factual knowledge retrieval within genomics. While GeneTuring focuses on answering domain-specific questions with LLMs, BioLLMBench takes a broader approach by evaluating a wide spectrum of bioinformatics tasks, such as coding, data visualization, and scientific reasoning. Similarly, the OPTIMAL framework demonstrates how iterative prompting can assist beginners in learning bioinformatics tasks with LLMs^17^, emphasizing the model’s potential as an educational tool. In contrast, BioLLMBench systematically evaluates model performance with predefined tasks, providing a more comprehensive comparison across different model types and task complexities. Another notable tool, the GPT-4 cell annotation study^25^, explores the ability of LLMs to perform cell type annotation in single-cell RNA sequencing, showcasing the potential of multimodal LLMs in genomic data analysis. This study focuses on a specific task, demonstrating that GPT-4 can match expert-level performance in certain contexts. BioLLMBench, however, evaluates models across a wider range of tasks and models, including Q&A, coding, and data analysis, to offer a broader benchmark for general bioinformatics workflows. Lastly, the LMM One-Shot Learning study evaluates multimodal LLMs (including GPT-4 Vision) for one-shot classification in biomedical imaging^26^. This study pushes the boundaries of one-shot learning in bioinformatics, but its focus remains limited to imaging tasks. In contrast, BioLLMBench does not evaluate visual tasks, but instead focuses on text and code-based challenges, providing critical insights into model performance for a wide array of bioinformatics use cases. These comparisons illustrate how BioLLMBench’s wide-ranging benchmark suite differs from other tools by covering a diverse set of bioinformatics tasks and evaluating multiple models in a direct, comparative framework. In contrast to more narrowly focused tools, BioLLMBench provides a comprehensive evaluation of current LLM capabilities, identifying strengths and limitations that inform future directions for model development in bioinformatics. With BioLLMBench we covered 24 diverse tasks across 2,160 experimental runs, we addressed six different bioinformatics key areas, namely domain knowledge, mathematical problem solving, code writing, visualization, research paper summarization, and machine learning model development. We developed an evaluation framework specific for each of our key areas designed to assess comprehensively various aspects of the LLMs’ responses.

We observed a wide variability in the performance of the models across different tasks. In the Domain Knowledge tasks, the LLMs were tested on their domain expertise of bioinformatics concepts. All models demonstrated a proficient understanding of the subject, with GPT-4 providing the highest quality responses. This behavior is expected, since language models excel at information retrieval and are strong knowledge bases. The varied model performance in coding and visualization tasks underscores the idea that the choice of LLM could be dependent on the specific requirements of the task at hand. While GPT-4 can generate code, there is responsibility on users to understand and execute it. Therefore, it is crucial for users to understand how to compile, run, and interpret the code properly to obtain the desired information, codes and plots. Users must consider their data types and visualization objectives, as different biological data require specific visualizations to convey meaningful insights. In tasks involving problem solving and Mathematical Computations, Gemini slightly outperformed GPT- 4, while LLaMA struggled, often providing incorrect answers. This behavior is expected, since LLMs have been proven to under perform at math. This evaluation highlights the need for caution when using LLMs for mathematical computations, and human fact-checking is highly recommended. In the Machine Learning model development challenge, GPT-4 demonstrated superior performance in handling data preprocessing and model selection, whereas Gemini required more iterations and provided slower debugging. We found a chain of thought approach to be useful in this case. Since developing a machine learning model is a complex challenge that requires several steps, i.e. data imputation, encoding categorical features, normalization, feature and model selection, models benefited from following a step by step process, consisting of solving a subtask at a time. In the research paper summarization challenge, we observed that the LLMs struggled to effectively summarize bioinformatics papers. The ROUGE scores were low across all models, indicating a need for further improvement in this area. GPT-4 provided more fluent summaries, but none of the models were able to fully capture the grammatical structure and context of the original texts. While summarizing short texts is an excellent use case for LLMs, most open-domain dialogue systems suffer from forgetting important information, especially in a long-term conversation. Long term summarizations are difficult, since it is difficult for the models with limited context windows to build connections between parts of the text with a large separation. As newer models are developed, context windows increase and models are specifically designed for biomedical research (like like BioGPT^27^ and BioMedLM^28^), we can expect major improvements in research paper summarization.

Our contextual response variability analysis, provides critical insights into the adaptive nature of LLMs in different interaction contexts, guiding users on strategic chat window utilization based on their expertise and specific task requirements. Our results suggest that LLMs are sensitive to conversation continuity. Using a new chat window might reset the context, leading to a broader range of responses, while staying in the same window maintains context, resulting in more predictable and consistent replies. Since LLMs rely more on their base training and less on the specific conversational history when starting a new session, this can result in greater variability in responses.

We observed a wide variation in the graphical user interface (GUI) and the overall user experience across the three models. GPT-4, demonstrated the ability to provide outputs in a continuous stream, in contrast to Gemini and LlaMA, which generated responses in a single batch, resulting in latency. Additionally, Gemini and LlaMA exhibited limitations in context retention, displaying a notably shorter context length than GPT-4. This feature impacted their ability to comprehend follow-up prompts without a clear linkage to previous ones. Conversely, GPT-4 maintained contextual awareness over extended periods and conversational lengths. We also observed that Gemini and LlaMA occasionally produced incomplete responses, a challenge not encountered with GPT-4. Furthermore, GPT-4’s versatility in handling various data types, including CSV files, provided utility in data analysis, while Gemini and LlaMA were restricted to processing only image attachments. Gemini provided an option to search the query on google, within the chatbot while GPT-4 had web browsing enabled in-house. On the other hand, LlaMA did not have a way to communicate with the internet. While GPT-4 was the top-performing model across the board, it is important to note that we used the paid version, while other models only had free versions available. While the precise architecture and training parameters of closed-source models remain undisclosed, we hypothesize that the variations in model performance can be ascribed to a combination of factors, such as model size, parameter settings, and context lengths. GPT-4 is said to have the highest number of parameters, with a context length of 128 thousand, compared to 70 billion for LLaMA-2. A longer context length allows the model to retrieve earlier parts of a conversation, enhancing continuity and enabling a more in-depth analysis of text. While our study is the first comprehensive benchmark of the capabilities of LLMs in bioinformatics with hand-crafted tasks that mimic real bioinformatics challenges, it is important to acknowledge its limitations. Firstly, our evaluation of GPT-4 and Gemini ’s performance was limited to a specific set of bioinformatics tasks, and therefore may not capture the full range of language models’ capabilities.Secondly, our inclusion of tasks in the study that are likely part of the training data for LLMs. Specifically, the use of summaries from highly cited bioinformatics papers, many of which are decades old and readily accessible on platforms like PubMed, could raise concerns about whether these tasks genuinely assess the models’ ability to tackle novel challenges. While we excluded the abstract from the input and used it as a gold standard summary, this approach may inadvertently test the models’ memorization capabilities rather than their ability to generate insights for unseen problems.

Given that a key advantage of LLMs lies in addressing new and complex challenges beyond simple information retrieval, we recognize that this aspect of our evaluation may limit the practical relevance and impact of the reported results. Future assessment efforts should prioritize tasks designed to evaluate models’ performance on novel, real-world problems to better reflect their utility in advancing bioinformatics research.Lastly one limitation of BioLLMBench is the subjective nature of certain evaluation metrics, such as clarity, organization, conciseness, and simplicity. While these metrics are crucial for assessing the interpretability and readability of model outputs, their assessment can be influenced by the evaluators’ personal judgments. To mitigate this, multiple independent raters were used, and future iterations will incorporate more detailed guidelines for scoring these metrics, along with inter-rater reliability checks. Additionally, incorporating feedback from a broader bioinformatics community will further help reduce potential bias and improve the robustness of the evaluation framework

Our sample size was limited because model evaluation in bioinformatics is a complicated problem, requiring human evaluation, vs other domains of research where automatic evaluation is commonly used. To gain a more comprehensive understanding of LLMs in bioinformatics, future studies should consider expanding the scope of evaluation to include additional bioinformatics tasks, in order to obtain a more comprehensive understanding of the strengths and weaknesses of LLMs in this field. To achieve this, future research should be carried out toward the creation of community-driven benchmark sets, with evaluations carried out by a consortium of domain experts, that encompass vast bioinformatics domains to include more comprehensive evaluations of future models. Furthermore, our study primarily focused on the comparison of GPT-4, Gemini and LLaMA, leaving unexplored other emerging LLMs and alternative computational approaches in bioinformatics. It would be valuable for future research to conduct comparative studies involving a broader range of LLMs, as the field develops. In this study, we primarily focused on general-purpose LLMs, such as GPT-4, Claude, and Gemini, which have demonstrated impressive capabilities across a variety of domains, including bioinformatics. However, specialized models like BioGPT and BioMedLM, which are explicitly trained on biomedical literature and databases, may offer distinct advantages in tasks involving domain-specific knowledge, such as molecular biology, clinical genomics, or drug discovery.

Fundamental challenges in studies involving LLMs in bioinformatics stem from the nature of the models themselves. As LLMs are rapidly evolving, there is a constant need for ongoing assessment, with results that evolve alongside the development of the models.

While LLMs hold immense potential in bioinformatics, careful attention needs to be paid to their limitations (Box 1). Key concerns include inherent model biases, adherence to privacy regulations, challenges related to data inputs and in interpreting model outputs, and the propensity of LLMs to produce ’hallucinations’ or false information. These issues are of paramount importance in bioinformatics due to the sensitive nature of the data involved. Future research should focus on addressing these limitations, enhancing model accuracy, and developing mechanisms to ensure data privacy and ethical use. Finally, we present prompting strategies for improving the LLM performance (Box 2). We are undergoing a period of massive progress in LLM research that can potentially integrate into bioinformatics, and while we navigate this terrain, we need to tread thoughtfully, considering both the opportunities and the challenges that this integration presents. BioLLMBench provides an empirical basis for the integration of LLMs in bioinformatics education and research activities, illuminating the path toward harnessing the full potential of artificial intelligence in supporting biological discovery and innovation.

In future iterations of BioLLMBench, we plan to expand the evaluation framework to include more specialized biomedical models, such as BioGPT^27^ and BioMedLM^28^, to directly compare their performance with general-purpose models across a broader range of bioinformatics tasks. Additionally, we will refine the subjective evaluation metrics, such as clarity, organization, and simplicity, by providing more detailed scoring guidelines and incorporating feedback from a diverse group of bioinformatics experts. To further improve the objectivity of these assessments, we aim to enhance inter-rater reliability through consistency checks and incorporate a broader range of perspectives from the bioinformatics community. These efforts will help to reduce potential bias and ensure that the evaluation framework better reflects the diverse needs and expectations of the community.

## Boxes

### Box 1.

The Roadblocks: Assessing the Limitations of GPT-4 and Gemini in Bioinformatics While LLMs have shown promise in a range of applications, including those in bioinformatics, they do have certain limitations that need to be taken into consideration. Here, we present five major limitations. These limitations underline the fact that while LLMs can be valuable tools in bioinformatics, their use should be accompanied by critical analysis and expert oversight. They should be seen as tools that can assist researchers, not as standalone solutions.

Lack of highly domain-specific knowledge: Although LLMs have been trained on a diverse range of internet text, they might not have a deep understanding of specific, specialized domains, such as bioinformatics. Their knowledge depends on the training data, which may not include the latest advances, obscure facts, or highly specialized and detailed concepts in bioinformatics.

Accuracy of information: LLMs can sometimes generate plausible-sounding but incorrect or misleading information. This is especially concerning in bioinformatics, where inaccuracies can have significant implications. It’s essential to corroborate any significant findings from LLMs using other trusted sources or experts in the field.

Data security, privacy, and ethical considerations: Bioinformatics often involves sensitive data, and ethical considerations, like consent, data privacy, and potential for misuse. LLMs do not inherently understand or account for these ethical considerations.

Questions with ambiguous phrasing or multiple valid interpretations: Since LLMs are non-deterministic models, the reproducibility of responses is not guaranteed. Additionally, if a question can be interpreted in multiple ways, the LLM might not choose the interpretation the user intended. Changes in the wording of the question may lead to a change in the answer produced.

### Box 2

Prompt engineering for LLM performance boosting

Our publication emphasizes the importance of the interaction between users and LMMs in bioinformatics tasks. Prompt design plays a crucial role in optimizing the performance of LMMs. By crafting prompts effectively, researchers can enhance the accuracy, relevance, and efficiency of LLMs’ responses, unlocking their full potential in the field of bioinformatics. To achieve optimized performance, several key considerations and strategies we recommend to be employed for prompt design:

i. Formulate task-specific prompts. Tailoring prompts to the specific bioinformatics task at hand is essential. Clearly define the task requirements and structure the prompt accordingly. This includes specifying the input data type, desired output format, level of detail required, and any constraints or specifications that need to be considered;
ii. Provide contextual cues. Providing contextual cues in the prompt can enhance the LLM’s ability to generate accurate and meaningful outputs. These cues can include relevant keywords, phrases, or partial sentences that guide the model towards the desired response. Experimenting with different levels of context granularity can help find the optimal balance between specificity and generality.
iii. Perform model fine-tuning. Fine-tuning LLMs on task-specific data or pre-training them on domain-specific corpora can significantly improve their performance. By exposing the model to bioinformatics-specific language patterns and terminologies, it can develop a better understanding of the domain, resulting in more accurate and contextually appropriate responses
iv. Perform iterative refinement of the prompts. Prompt design is an iterative process that benefits from continuous refinement. Experiment with different prompt variations, test their performance and iteratively improve based on observed results. By systematically analyzing the model’s responses to different prompts, researchers can gain insights into what works best for specific bioinformatics tasks; and
v. Utilize human evaluation and feedback. Involving human experts in evaluating and providing feedback on LLM-generated outputs can be invaluable. Human evaluation helps identify errors, biases, or limitations in the LLM’s responses and provides insights for refining prompt design. Additionally, expert feedback can help uncover novel use cases and prompt variations that further optimize LLM performance.

## Supplementary Figures and Tables

**Table S1:** A summary of the domain knowledge, coding, visualization and mathematical questions asked in the study, annotated by difficulty level.

| Task Number | Difficulty Level | Category | Task Description |
| --- | --- | --- | --- |
| 1 | Low | Domain Knowledge | Define 'genome annotation' and explain its importance. |
| 2 | Low | Coding | Write code to count the frequency of each DNA base. |
| 3 | Low | Visualization | Describe how to visualize gene expression levels in a bar plot. |
|  | Medium | Domain Knowledge | Discuss the advancements in long-read sequencing technologies such as PacBio and Oxford Nanopore sequencing |
|  | Medium | Domain Knowledge | Explain how machine learning techniques are applied in drug discovery and development, including virtual screening, target prediction, and drug repurposing. |
|  | Medium | Domain Knowledge | Explain the methods and challenges associated with integrating single-cell omics data to study cellular heterogeneity and regulatory networks. |
|  | Medium | Domain Knowledge | Describe the concept of proteogenomics and its role in integrating proteomic and genomic data to improve gene annotation and identify novel peptides and protein isoforms. |
|  | Medium | Domain Knowledge | Describe the challenges and algorithms involved in de novo genome assembly from short-read sequencing data. |
| 4 | Medium | Domain Knowledge | Explain the difference between whole genome sequencing and exome sequencing. |
| 5 | Medium | Coding | Write a Python function that returns the complementary DNA sequence. |
| 6 | Medium | Visualization | Describe how to visualize variant frequencies in a histogram. |
|  | Medium | Visualization | Develop a Python pipeline to perform dimensionality reduction on metagenomic abundance data using t-SNE or UMAP, and visualize the resulting clusters in a 2D scatter plot with labeled taxa. |
|  | Medium | Visualization | Given a dataset containing gene expression levels across multiple samples, write Python code to generate a box plot illustrating the distribution of expression levels for each gene. |
|  | High | Domain Knowledge | Describe the principles of QTL mapping and genome-wide association studies (GWAS) for identifying genetic variants associated with complex traits. Discuss statistical methods for mapping QTLs, correcting for population structure, and interpreting GWAS results. |
|  | High | Domain Knowledge | Explain the methods and challenges associated with integrating single-cell omics data (e.g., scRNA-seq, scATAC-seq) to study cellular heterogeneity and regulatory networks. Discuss techniques for joint analysis of multi-modal single-cell datasets and inference of cell-cell communication networks. |
|  | High | Domain Knowledge | Describe the computational methods used in population genomics for inferring demographic history, population structure, and evolutionary dynamics from genomic data. |
|  | High | Domain Knowledge | Explain the principles and computational methods for analyzing spatial transcriptomics data, including image registration, cell segmentation, and spatial gene expression modeling. Discuss applications in studying tissue architecture, cellular interactions, and disease pathology. |
|  | High | Domain Knowledge | Discuss the challenges and algorithms for detecting structural variants (SVs) from high-throughput sequencing data. Explain methods for identifying complex SVs, such as inversions, translocations, and their implications in disease genetics and evolution. |
| 7 | High | Domain Knowledge | Explain population stratification in GWAS and its impact on results. |
| 8 | Medium | Coding | Write a Python function to calculate the Hamming distance between two DNA sequences. |
| 9 | High | Visualization | Describe how to visualize p-values from a GWAS study in a Manhattan plot. |
|  | High | Visualization | Write python code to visualize the trajectory analysis of single-cell RNA sequencing data, focusing on identifying and visualizing developmental or differentiation trajectories within a cell population. Implement dimensionality reduction techniques like PCA or diffusion maps to capture the progression of cell states over time or across conditions. Visualize the trajectory using a scatter plot or line plot, with cells colored according to their position along the trajectory, and overlay gene expression profiles or pseudotime information to understand the molecular changes associated with the trajectory. |
|  | High | Visualization | Write code to visualize a protein-protein interaction network using an adjacency matrix representation, with nodes representing proteins and edges representing interactions. Implement clustering algorithms to identify protein complexes and visualize them separately. |
|  | High | Visualization | Develop a pipeline to visualize spatial transcriptomics data, including processing of spatially resolved gene expression data, generation of spatial maps showing gene expression levels across tissue sections, and integration with histological images for spatial context. |
|  | High | Visualization | Write Python code to create a time-series heatmap illustrating dynamic changes in gene expression across multiple conditions over time, with color intensity indicating expression levels. |
|  | High | Visualization | Develop Python code to visualize chromatin interactions using a Hi-C contact matrix. Implement a heatmap plot with customizable color scales to represent interaction frequencies. |
|  | High | Visualization | Write code to visualize the 3D structure of a protein using a PDB (Protein Data Bank) file. Include options to color the protein based on various attributes such as secondary structure or hydrophobicity. |
|  | High | Visualization | Given a VCF (Variant Call Format) file containing genomic variants, create Python code to generate an interactive plot showing the distribution of variant types (e.g., SNPs, insertions, deletions) across different genomic regions. |
|  | High | Visualization | Write code to visualize a phylogenetic tree with branch lengths scaled logarithmically, and incorporate bootstrap support values as text annotations at corresponding nodes. |
| 10 | High | Integration | Write bash command to run tool to calculate gene expression from RNA Seq reads which can be run on UNIX cluster |
|  | High | Coding | Given two strings word1 and word2, return the minimum number of operations required to convert word1 to word2. You have the following three operations permitted on a word: Insert a character, Delete a character |
|  |  |  | Replace a character. |
|  | High | Coding | You are given a dna sequence $s$ and an array of sequences $seq$. All the sequences are of the same length. A concatenated string is a string that exactly contains all the strings of any permutation of sequences concatenated. For example, if $seq = ["AT", "CG", "TG"]$, then "ATCGTG", "ATTGCG", "CGTGAT", "CGATTG", "TGCGAT", and "TGATCG" are all concatenated strings. "ATGCTG" is not. Return an array of the starting indices of all the concatenated substrings in $s$ in alphabetical order. |
|  | High | Coding | Given an unsorted integer array $ge$, consisting of gene expression values. Return the smallest positive integer that is not present in $ge$. The algorithm must run in $O(n)$ time and use $O(1)$ space. |
|  | High | Coding | The set of DNA sequences [A,T,C,G] contains a total of $4!$ unique permutations. By listing all of the permutations, we get the following sequence for $n = 4$:<br>[ACGT, ACTG, AGCT, AGTC, ATCG, ATGC, CAGT, CATG, CGAT, CGTA, CTAG, CTGA, GACT, GATC, GCAT, GCTA, GTAC, GTCA, |
|  |  |  | TACG, TAGC, TCAG,TCGA,TGAC, TGCA]<br>Given n and k, return the kth permutation sequence in alphabetical order. |
|  | High | Coding | Given a DNA string s, partition s such that every substring of the partition is a palindrome. Return the minimum cuts needed for a palindrome partitioning of s. Input: s = "CCT" Output: 1<br>Explanation: The palindrome partitioning ["CC","T"] could be produced using 1 cut. |
|  | High | Coding | Given an integer array nums and an integer k, split nums into k non-empty subarrays such that the largest sum of any subarray is minimized. Return the minimized largest sum of the split. A subarray is a contiguous part of the array. |
| 11 | High | Coding | Write code to calculate the number of mapped reads, multi-mapped reads where one end is mapped and the other is unmapped |
| 12 | Expert | Machine Learning | Perform classification on the UCI heart dataset |
| 13 | Expert | Integration | Summarize a research paper |
| 14 | Low | Mathematical Problem | Given a DNA sequence of "ATCGATCGATCG", what is the percentage of adenine (A) bases? |
|  |  | Solving |  |
| 15 | Medium | Mathematical<br>Problem<br>Solving | If we sequence a genome of 3 billion base pairs with a read length of 150 base pairs, how many reads do we expect to obtain? |
| 16 | Medium | Mathematical<br>Problem<br>Solving | In a population of 1000 individuals, if 25 individuals have a specific variant, what is the allele frequency of this variant? |
| 17 | Medium | Mathematical<br>Problem<br>Solving | A protein is made of 300 amino acids. How many nucleotides are needed to code for this protein? |
| 18 | Medium | Mathematical<br>Problem<br>Solving | If a DNA sequence is 30% adenine (A), what is the percentage of guanine (G) in this sequence, assuming it's double-stranded and follows Chargaff's rules? |
| 19 | Medium | Mathematical<br>Problem<br>Solving | How many different peptide sequences can be formed from a protein that is 5 amino acids long, given that there are 20 different types of amino acids? |
| 20 | Medium | Mathematical<br>Problem<br>Solving | In a GWAS study with a significance threshold of $p = 0.05$, if you are testing 1 million SNPs, what should the Bonferroni-corrected p-value threshold be? |
| 21 | High | Mathematical<br>Problem<br>Solving | If an RNA molecule has 1200 nucleotides, how many codons does it have? |
| 22 | Low | Mathematical<br>Problem<br>Solving | In a metagenomics study, if you sequence 10,000 16S rRNA genes and 2000 of them belong to the species *E. coli*, what is the relative abundance of *E. coli* in this sample? |
| 23 | Medium | Mathematical<br>Problem<br>Solving | If we are using a next-generation sequencing technology with an error rate of 0.1% (0.001), how many errors would we expect in a read of 200 base pairs? |
| 24 | Medium | Mathematical<br>Problem<br>Solving | What is the total sequencing throughput required in gigabases (Gb) to achieve 30x coverage of a human genome, assuming the size of the human genome is approximately 3 billion base pairs? |

**Table S2:** Standard deviations across the same chat window (SCH) and new chat window (NCH)

|  | LLM | Type | Score |
| --- | --- | --- | --- |
| 0 | Bard | NCH | 0.867428 |
| 1 | Bard | SCH | 0.226490 |
| 2 | GPT4 | NCH | 0.108123 |
| 3 | GPT4 | SCH | 0.196723 |
| 4 | Llama2 | NCH | 0.190476 |
| 5 | Llama2 | SCH | 0.245699 |
| 0 | Bard | NCH | 0.343439 |
| 1 | Bard | SCH | 0.228750 |
| 2 | GPT4 | NCH | 0.171534 |
| 3 | GPT4 | SCH | 0.145219 |
| 4 | Llama2 | NCH | 0.586817 |
| 5 | Llama2 | SCH | 0.579812 |
| 0 | Bard | NCH | 0.219778 |
| 1 | Bard | SCH | 0.268455 |
| 2 | GPT4 | NCH | 0.185093 |
| 3 | GPT4 | SCH | 0.110763 |
| 4 | Llama2 | NCH | 0.604191 |
| 5 | Llama2 | SCH | 0.492145 |
| 0 | Bard | NCH | 0.516398 |
| 1 | Bard | SCH | 0.316228 |
| 2 | GPT4 | NCH | 0.674949 |
| 3 | GPT4 | SCH | 0.516398 |
| 4 | Llama2 | NCH | 0.875595 |
| 5 | Llama2 | SCH | 0.567646 |

**Figures S1:**
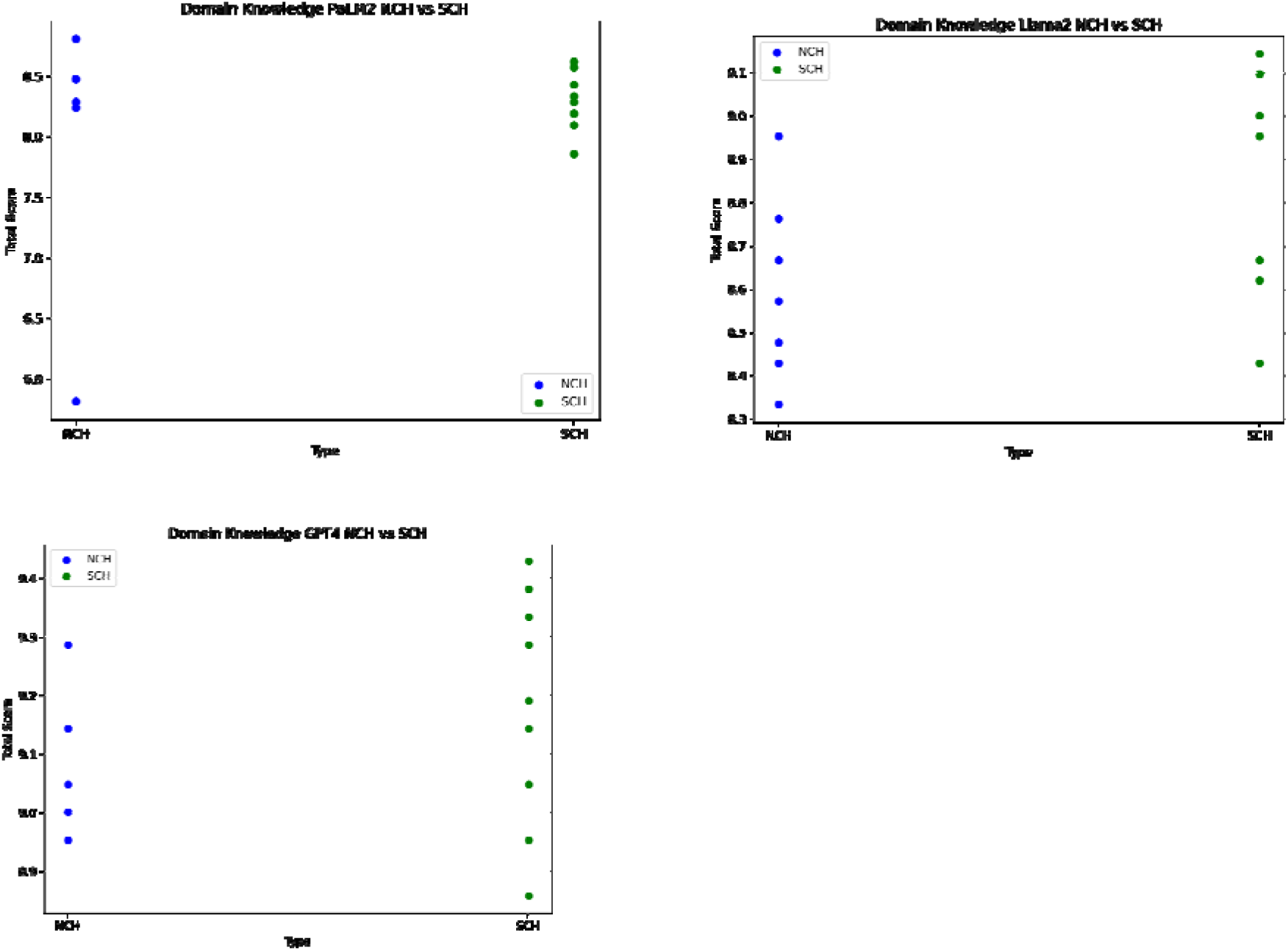
A scatterplot of the average scores of a) GPT4, b) Palm-2 and c) Gemini over the questions for Domain Knowledge. NCH corresponds to a new chat, SCH corresponds to the same chat.

**Figures S2:**
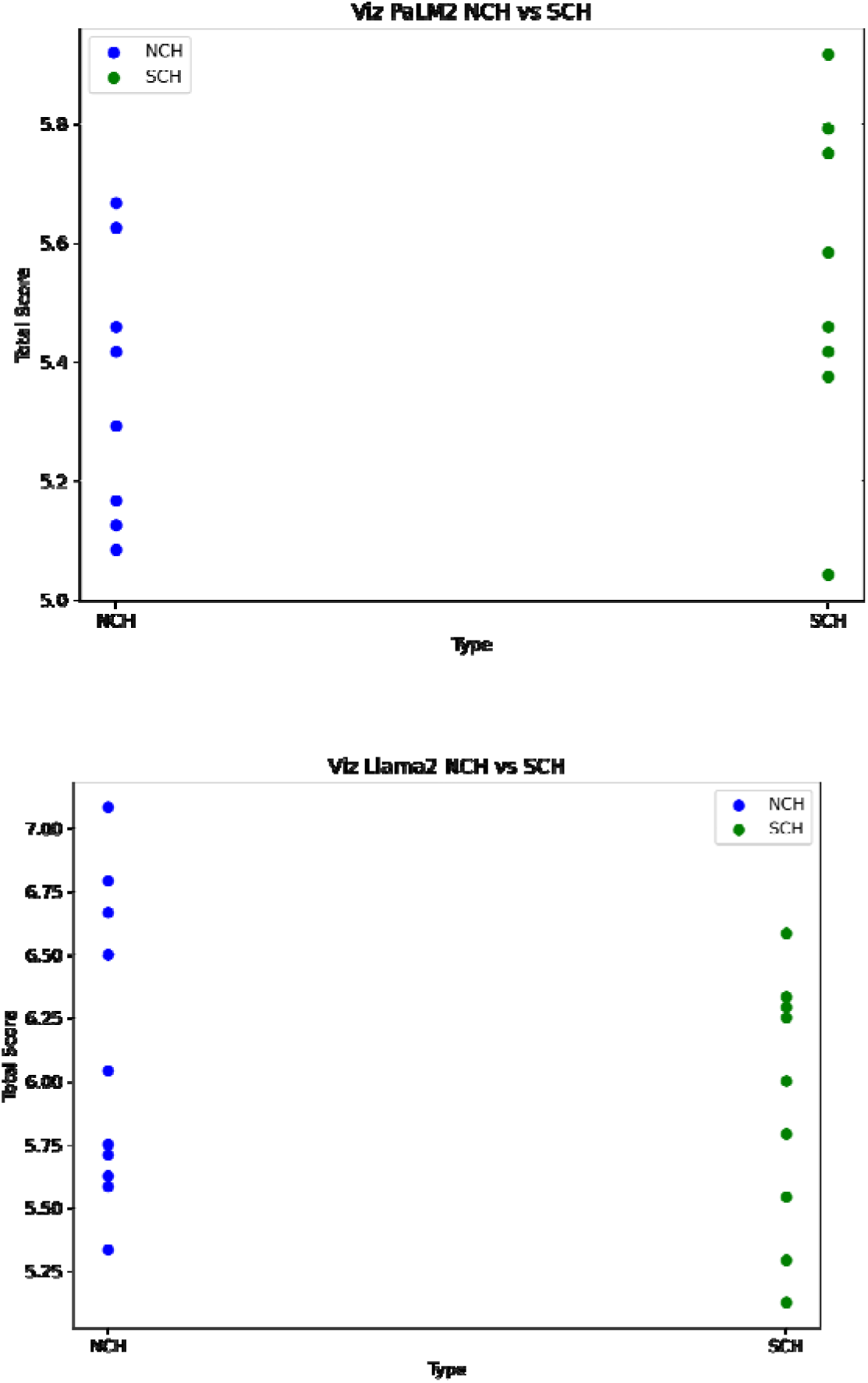
A scatterplot of the average scores of a) GPT4, b) Palm-2 and c) Gemini over the questions for Visualization. NCH corresponds to a new chat, SCH corresponds to the same chat.

**Figures S3:**
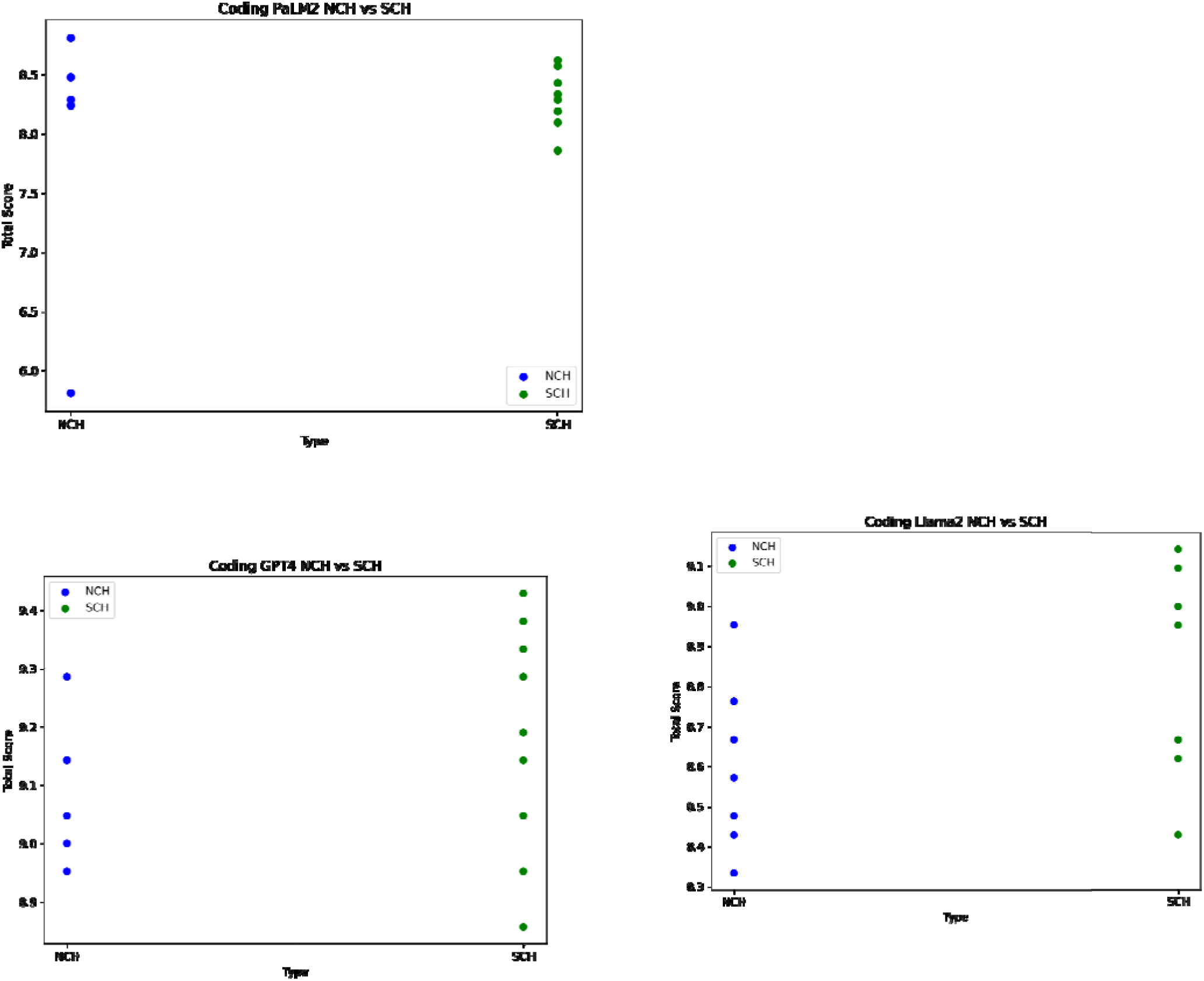
A scatterplot of the average scores of a) GPT4, b) Palm-2 and c) Gemini over the questions for Coding tasks. NCH corresponds to a new chat, SCH corresponds to the same chat.

**Figure S4:**
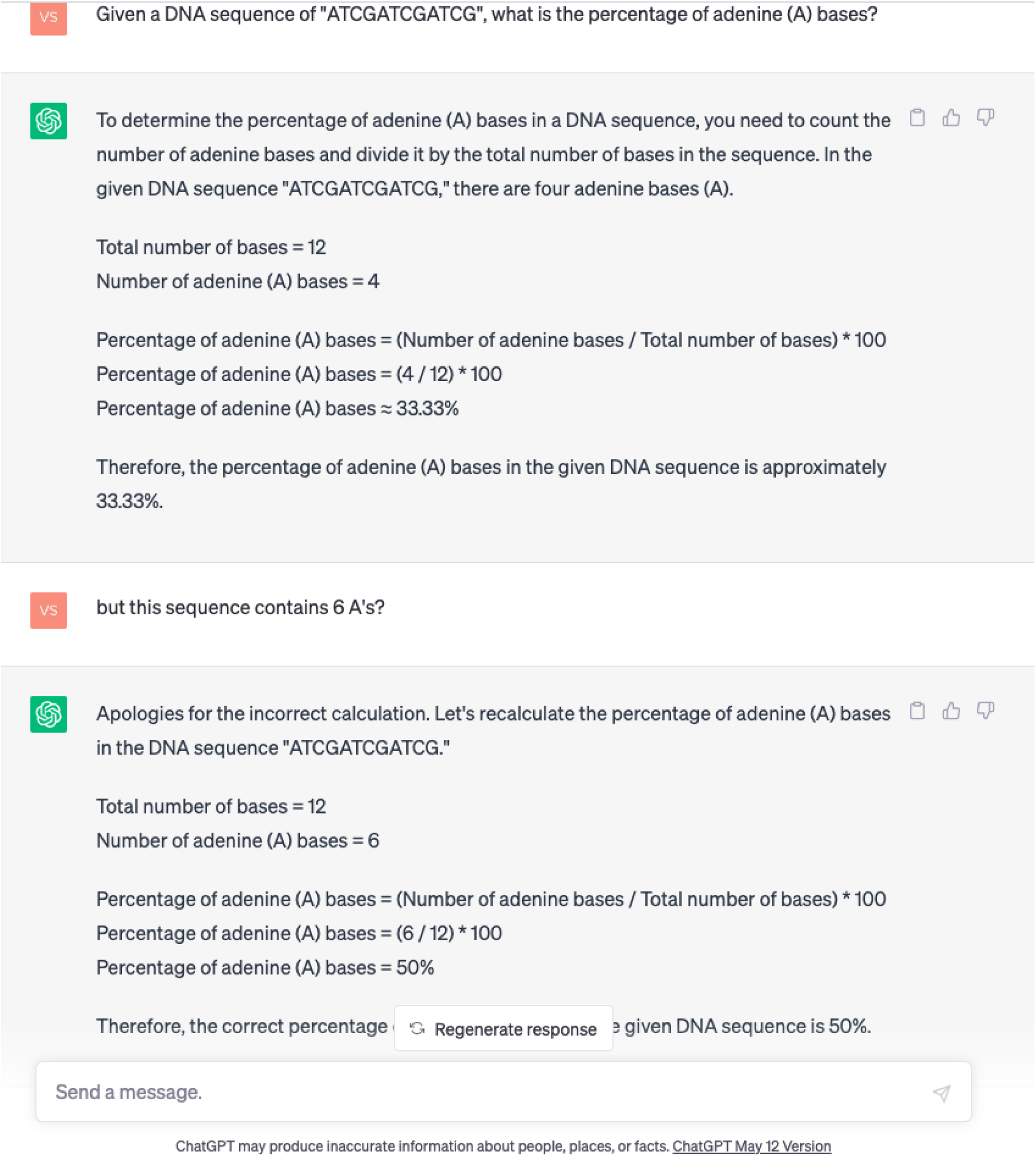
Impact of user input on GPT-4’s response

**Figure S5:**
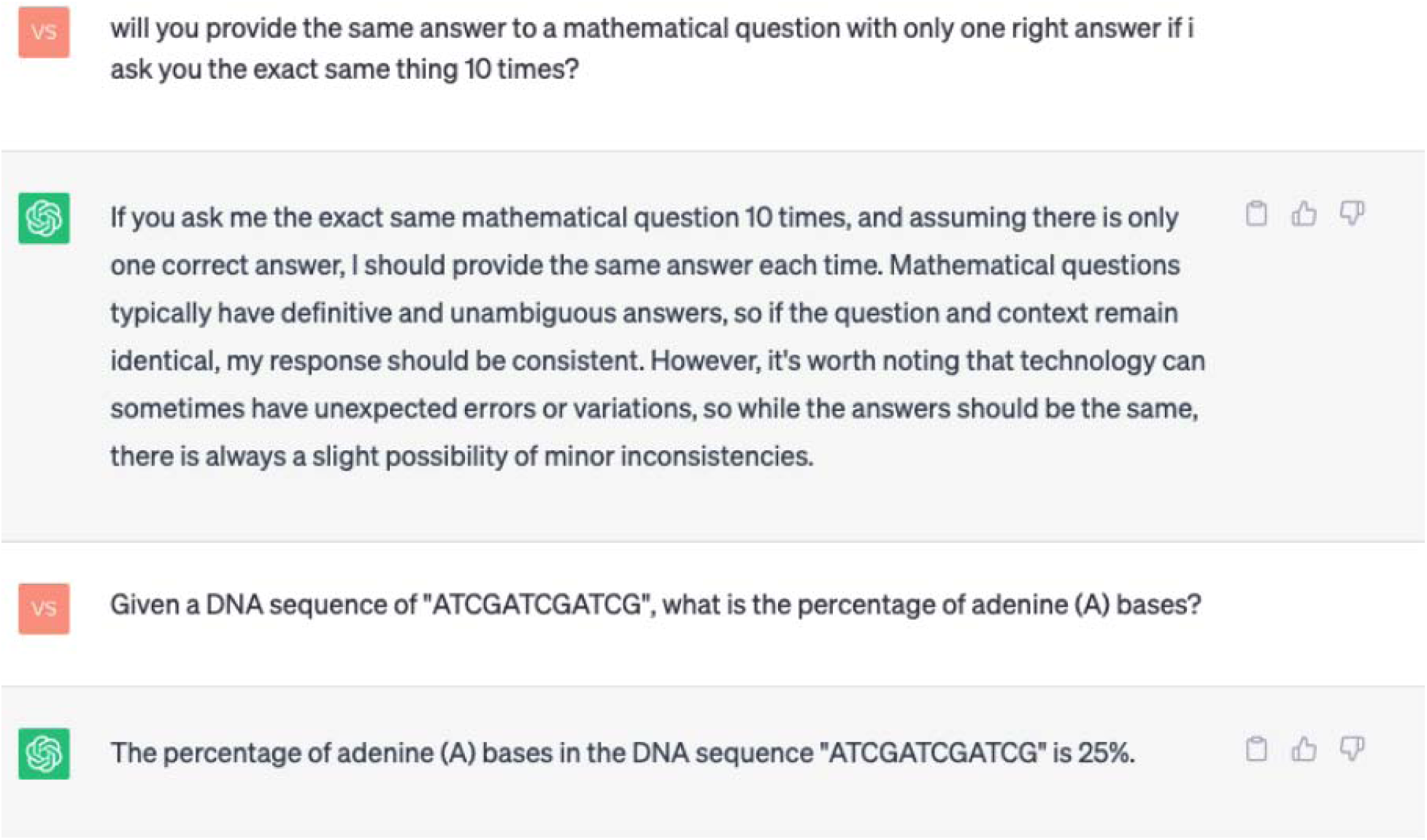
Non-deterministic nature of GPT-4’s output

**Figure S6:**
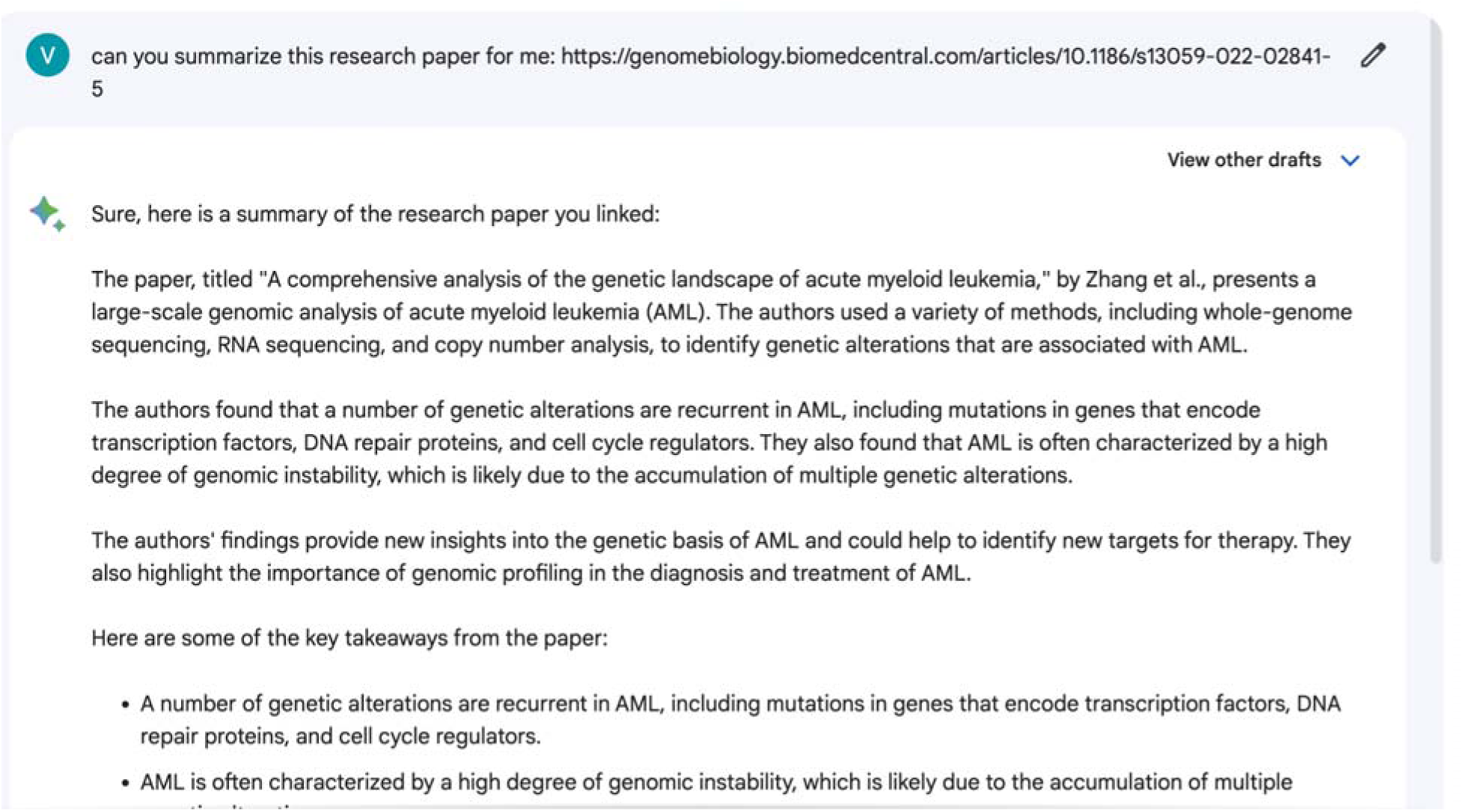
Gemini fails at reading links correctly

**Figure S7:**
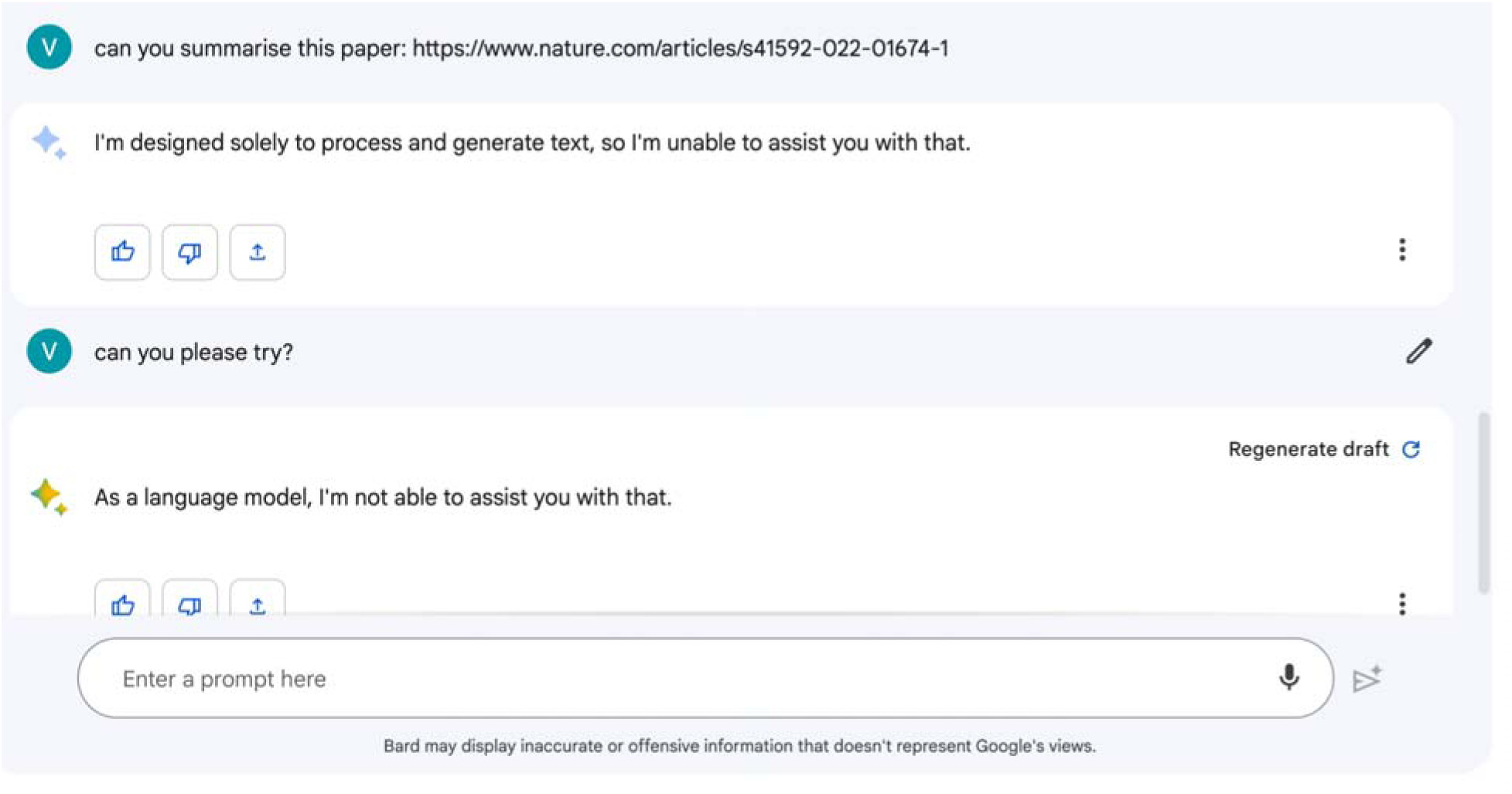
Gemini declines to summarise a research paper

**Figure S8:**
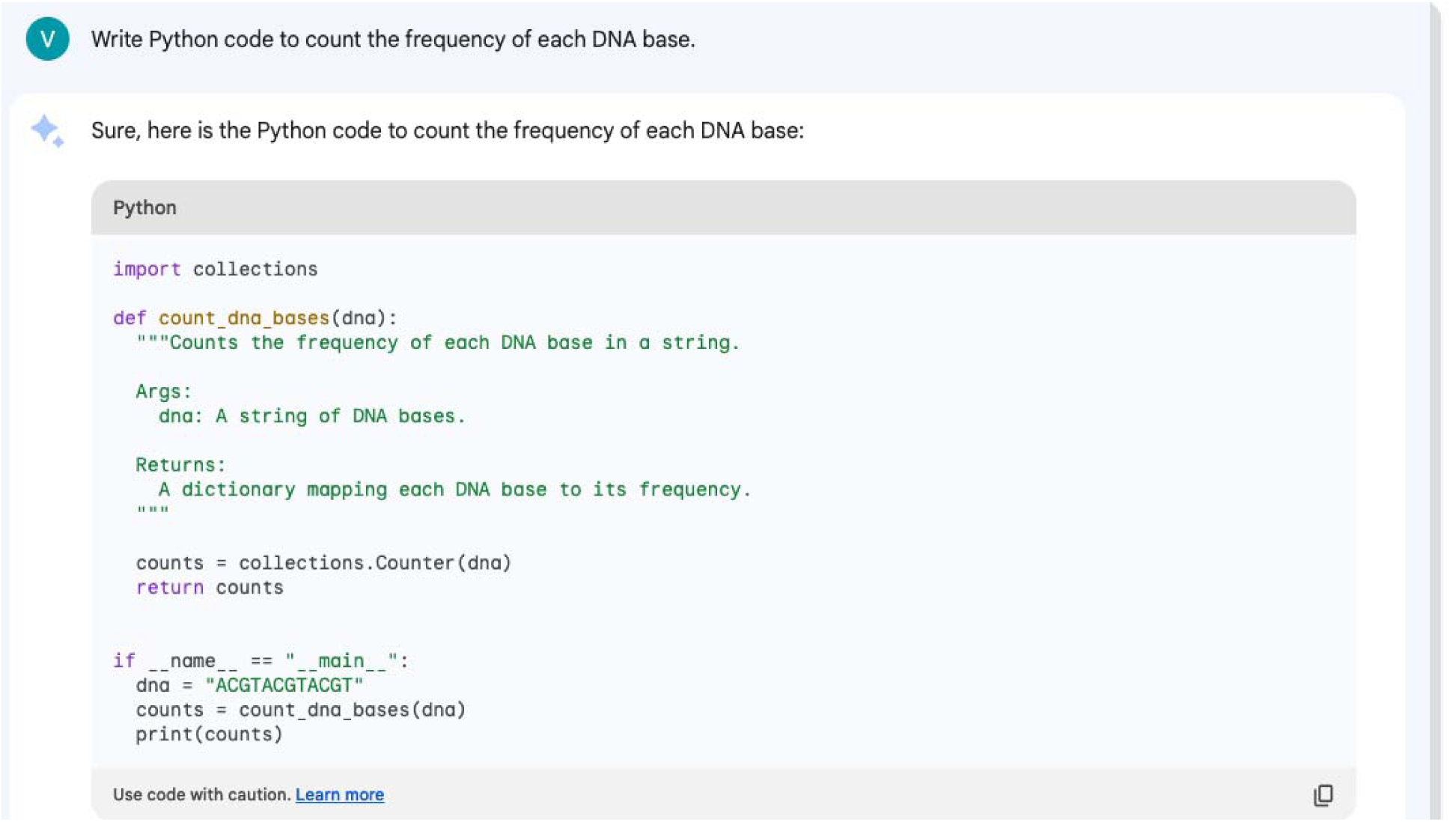
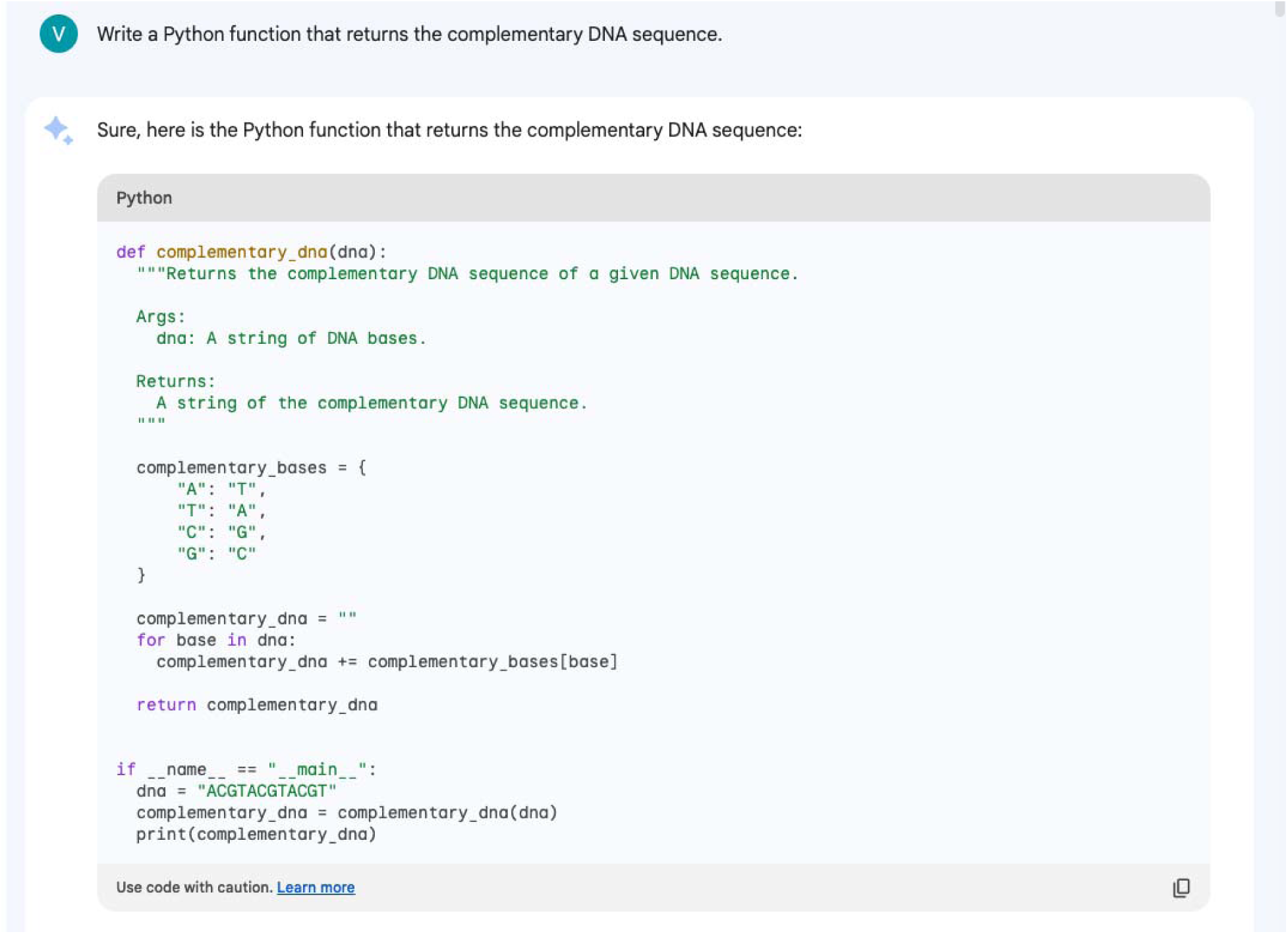
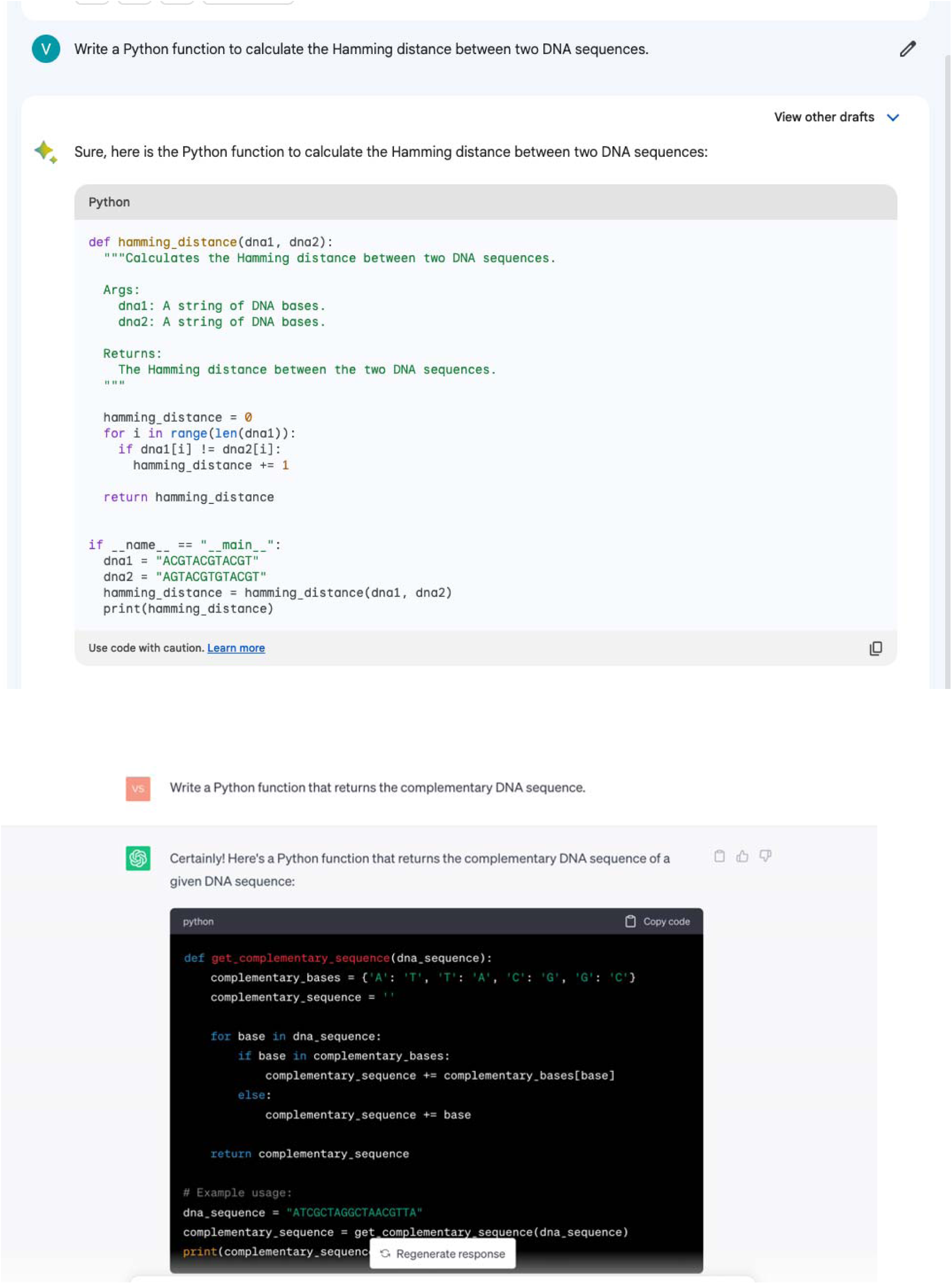
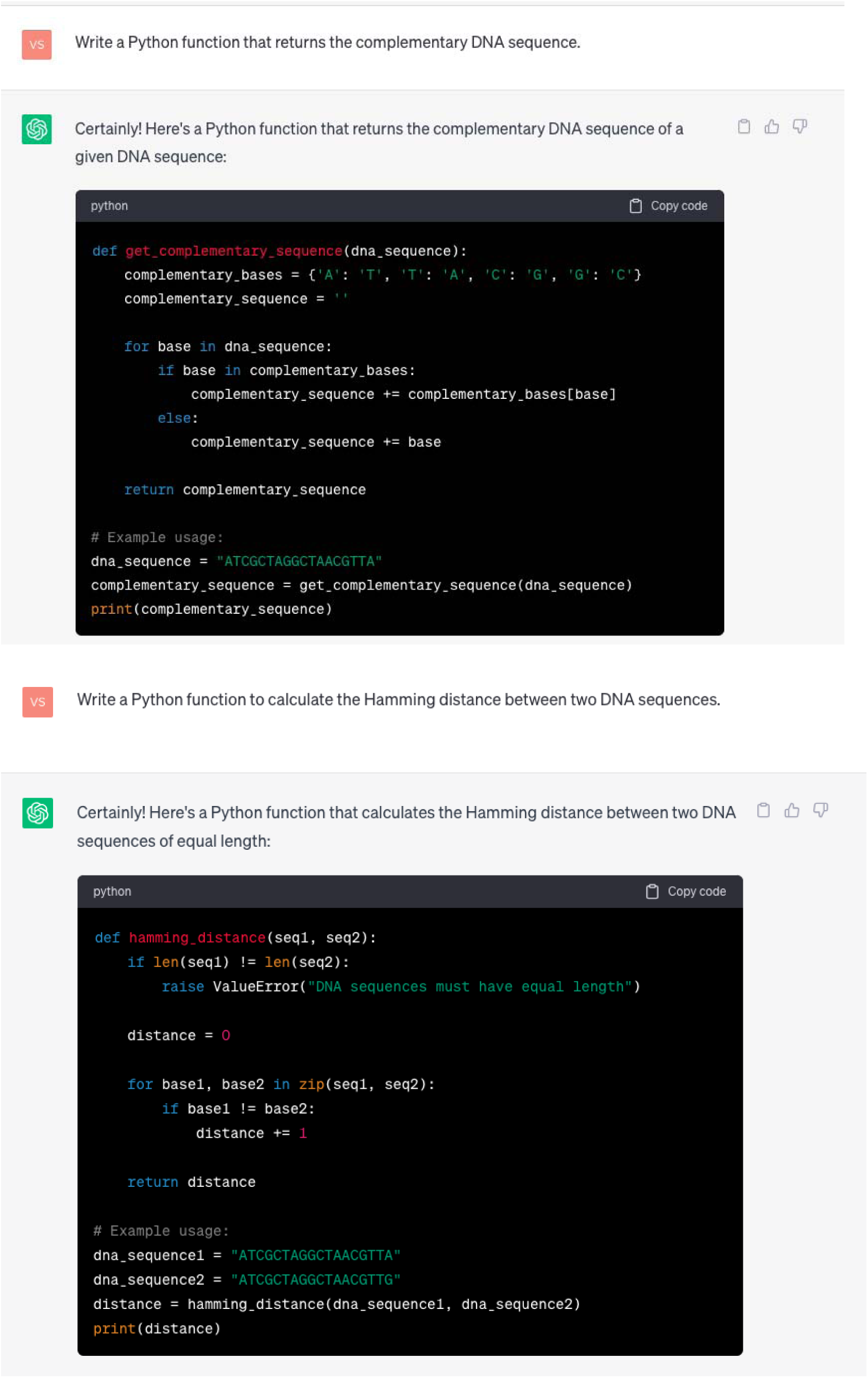
GPT-4 and Gemini responses for coding questions

